# A structural dendrogram of the actinobacteriophage major capsid proteins provides important structural insights into the evolution of capsid stability

**DOI:** 10.1101/2022.09.09.507160

**Authors:** Jennifer M. Podgorski, Krista Freeman, Sophia Gosselin, Alexis Huet, James F. Conway, Mary Bird, John Grecco, Shreya Patel, Deborah Jacobs-Sera, Graham Hatfull, Johann Peter Gogarten, Janne Ravantti, Simon White

**Affiliations:** Biology/Physics Building, Department of Molecular and Cell Biology, University of Connecticut, 91 North Eagleville Road, Unit-3125. Storrs, CT 06269-3125, USA; Clapp Hall, Department of Biological Sciences, University of Pittsburgh, 4249 Fifth Avenue, Pittsburgh, PA 15260, USA; Department of Structural Biology, University of Pittsburgh School of Medicine, Pittsburgh, PA, USA.; Institute for Systems Genomics, University of Connecticut, Storrs, CT 06268-3125, USA.; University of Helsinki, Molecular and Integrative Biosciences Research Programme, Helsinki, Finland.

## Abstract

Many double-stranded DNA viruses, including tailed bacteriophages (phages) and herpesviruses, use the HK97-fold in their major capsid protein to make the capsomers of the icosahedral viral capsid. Following the genome packaging at near-crystalline densities, the capsid is subjected to a major expansion and stabilization step that allows it to withstand environmental stresses and internal high pressure. Several different mechanisms for stabilizing the capsid have been structurally characterized, but how these mechanisms have evolved is still not understood. Using cryo-EM structure determination, structural comparisons, phylogenetic analyses, and Alphafold predictions, we have constructed a detailed structural dendrogram describing the evolution of capsid structural stability within the actinobacteriophages. The cryo-EM reconstructions of ten capsids solved to resolutions between 2.2 and 4 Ångstroms revealed that eight of them exhibit major capsid proteins that are linked by a covalent cross-linking (isopeptide bond) between subunits that was first described in the HK97 phage. Those covalent interactions ultimately lead to the formation of mutually interlinked capsomers that has been compared to the structure of chain mail. However, three of the closely related phages do not exhibit such an isopeptide bond as demonstrated by both our cryo-EM maps and the lack of the required residue. This work raises questions about the importance of previously described capsid stabilization mechanisms.

## Introduction

The HK97-fold (Figure 1 A) is ubiquitous in the biosphere and has been identified in viruses that infect the three domains of life^1,2,3^, as well as encapsulins^4^: protein shells used by bacteria for gene transfer and reaction confinement^5^. It is found across the Caudovirales order (the double-stranded DNA tailed phages), which is one of the largest groups of viruses in the biosphere and plays major roles in bacterial evolution and in carbon/nitrogen/phosphorus cycling^6^. Actinobacteriophages (bacteriophages infecting actinobacterial hosts) have been intensively studied with over 20,000 individual isolates, the vast majority of which are dsDNA phages. These phages are the central focus of integrated research-education programs^7, 8^, have provided tools for *Mycobacterium* genetics^9^, and show promise as therapies for drug-resistant *Mycobacterium* infections^10, 11^.

**Figure 1.**
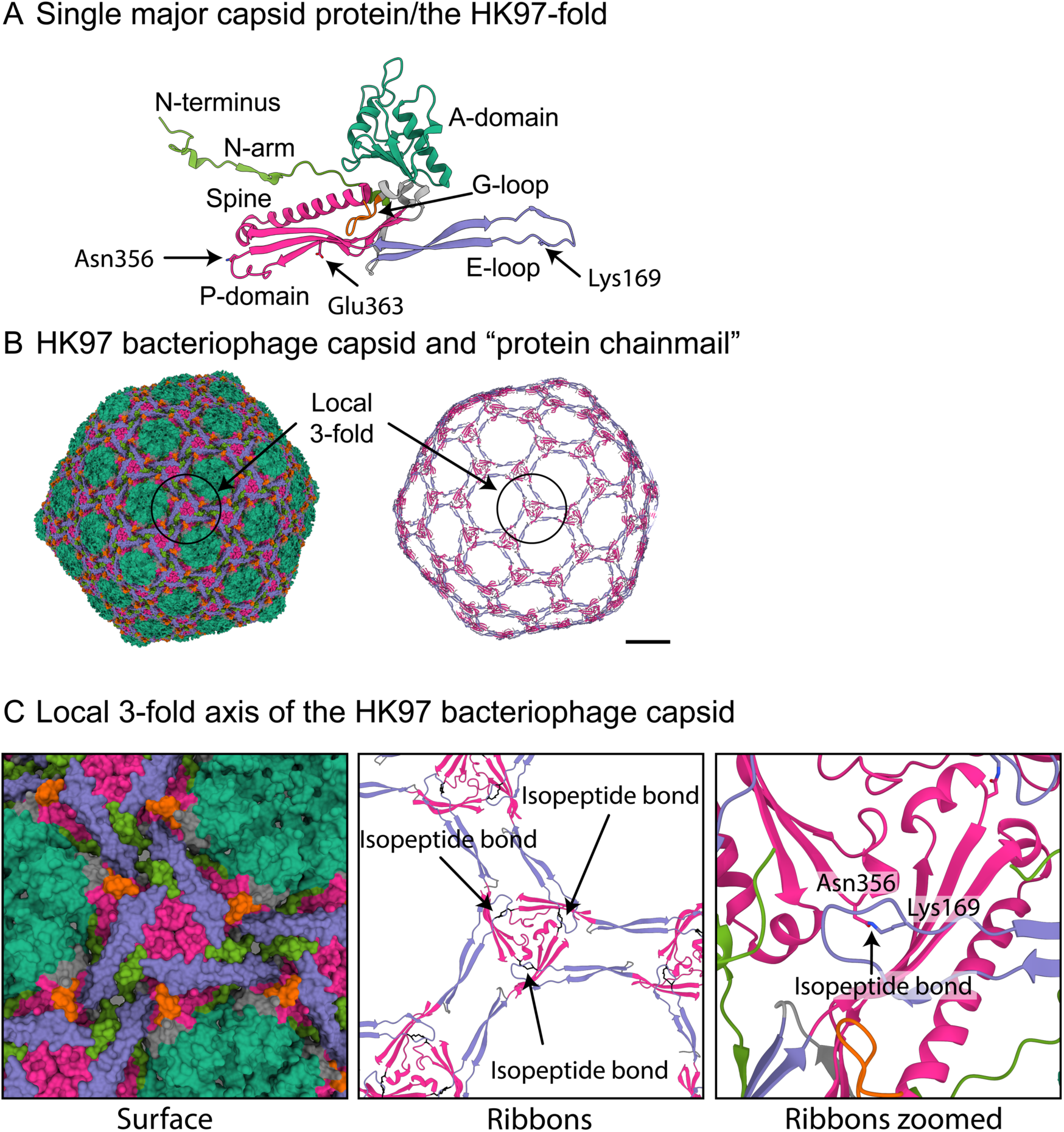
The HK97 bacteriophage: the prototypical HK97-fold. A) A single major capsid protein subunit from the HK97 bacteriophage showing the conserved domains of the HK97-fold: A-domain (teal), E-loop (purple), G-loop (orange), P-domain (magenta), and N-arm (green). The two amino acids (Lysine169 and Asparagine356) of the isopeptide bond are shown, as is Glu363, which catalyzes the bond formation. B) Model of the mature HK97 bacteriophage (1OHG) on the left and on the right a ribbon diagram highlighting the concatenated rings that are formed by the isopeptide bond. Coloring is the same as in A. Scalebar = 10 nm. C) A zoomed-in image of B showing the local 3-fold axis where the isopeptide bond is formed and where many stabilizing interactions in other bacteriophage capsids have been described.

Most of the major capsid proteins in the tailed phages use the HK97-fold (Figure 1 A) as the foundational block to build the capsid (Figure 1 B). To date, all structurally characterized HK97-folds have several conserved domains^12, 13^. They include the A domain (Figure 1A, teal) that forms the central core of the hexamer and pentamer capsomers of the capsid; the P-domain (Figure 1A, magenta) where the long spine helix is located; as well as the E-loop (Figure 1A, purple) and N-arm (Figure 1A, green). The E-loop and N-arm make long-range contacts with other major capsid proteins and play important roles in capsid stability^14^. The viral major capsid protein that uses the HK97-fold is assembled into an icosahedral shell consisting of eleven pentamers and different numbers of hexamers of the major capsid protein depending on the size and shape of the capsid. The icosahedral capsid is described by a T number that defines the number of major capsid proteins in the icosahedral shell - equal to the T number multiplied by sixty^15^. Within the dsDNA tailed phages, one pentamer is replaced by the portal to which the tail is bound and through which the DNA is packaged and then released.

The stability of the mature capsid is a key factor in the evolutionary success of phages^16^. The capsid needs to withstand various environmental conditions, and the pressure of the packed DNA genome^17,18,19^. The local 3-fold axes for each capsomer, where the P-domains of the pentamer and hexamer capsomers intersect, are thought to be important for capsid stability^16^ (Figure 1 B and C highlights one such 3-fold axis in the HK97-fold). The tailed phages use several different mechanisms to stabilize the capsid local 3-fold axes. The most common is a minor capsid protein, or ‘cement’, that binds to the local 3-fold axis and makes several contacts with the surrounding major capsid subunits^20, 21^. Others use catenated rings, with either non-covalent^22^ or covalent bonding^23^ mechanisms, that connect the major capsid proteins around the local 3-fold axis^14^. The major capsid protein of the HK97 phage (in which the HK97-fold was first characterized) uses a covalent isopeptide bond to cross-link a conserved asparagine (Asn356) in the P-domain of one major capsid protein with a conserved lysine (Lys169) in an adjacent E-loop of a different major capsid protein^23^ (Figure 1 B and C). The cross-linking is catalyzed by a nearby glutamic acid (Glu363) on a third major capsid protein subunit and aided by three other amino acids that form a hydrophobic pocket^24^. The cross-linking of all the major capsid proteins results in the catenated rings or “protein chainmail” that stabilizes the capsid around the local 3-fold axes (Figure 1 C).

Some phages have been characterized, for example, T5^25^, T7^26^, and phiRSA1^27^, that rely solely on intracapsomeric interactions and do not use cement or non-covalent/covalent bonding mechanisms. These different mechanisms of capsid stabilization make the HK97-fold highly adaptable and able to survive a wide variety of environments, from soil to hot springs and allows for the formation of structurally very diverse capsids that range in size from relatively small 50 nm diameter capsids^28,29,30^ to hundreds of nanometers in diameter “giant” capsids^31, 32^.

High-resolution structures of over twenty-five tailed phage capsids ^21–50^, viruses that infect archea^2^, and the human pathogenic Herpesvirus^3^ show that the HK97-fold is well conserved, even among viral capsids sharing little or no amino acid sequence similarity and using several different capsid stability mechanisms. However, these structures are from viruses that infect diverse hosts across all three domains of life and are so divergent from one another that only limited conclusions can be made about their evolution. We, therefore, carried out a systematic investigation of closely related phages infecting actinobacterial hosts to understand how capsid stability mechanisms are conserved and how they have evolved.

## Materials and Methods

### Production and purification of Phages for Cryo-Electron Microscopy

Phages were produced as previously described^51^. Briefly, twenty webbed plates were made for each phage with their host (Supporting Information Table 1) in top agar on Luria agar plates and incubated overnight at the temperatures shown in Supporting Information Table 1. Phages were extracted from the webbed plates using 5 mL of Phage Buffer (10 mM Tris-HCl pH 7.5, 10 mM MgSO_4_, 68 mM NaCl, 1 mM CaCl_2_) and incubated overnight at room temperature to allow diffusion of the phages into the Phage Buffer. The lysate was aspirated from the plates and centrifuged at 12,000× *g* for 15 min at 4 °C to remove cell debris. Phage particles were then pelleted using an SW41Ti swinging bucket rotor (Beckman Coulter, Brea, CA) at 30,000 rpm for 3 hours using 12.5 mL open-top poly clear tubes (Seton Scientific, Petaluma, CA). The phage particles in the pellet were then resuspended in 7 mL of Phage Buffer by gentle rocking overnight at 4 °C. The new phage lysate was subjected to isopycnic centrifugation with the addition of 5.25 g of CsCl to the 7 mL of phage lysate. The CsCl/phage solutions were centrifuged at 40,000 rpm in an S50-ST swinging bucket rotor (Thermofisher Scientific, Waltham, MA) for 16 h and the phage particle band (that appeared roughly halfway down the tube) was removed via side puncture with a syringe and needle. Phage particles were then dialyzed three times against Phage Buffer to remove CsCl. To do this the ∼1 mL of purified phages was placed into a Tube-O-Dialyzer Micro (G-Biosciences, St Louis, MO) with a 50 kDa molecular weight cut-off. The phages were then concentrated a final time by pelleting them at 75000 rpm in an S120-AT2 fixed angle rotor (Beckman Coulter, Brea, CA). The phage particles were then resuspended in 20 µL of Phage Buffer with gentle pipetting.

### Preparation of Cryo-Electron Microscopy Grids

Five microliters of concentrated phage particles (approximately 10 mg/mL) were added to Au-flat 2/2 (2 µm hole, 2 µm space) cryo-electron microscopy grid (Protochips, Morrisville, NC, USA) using a Vitrobot Mk IV (Thermo Fisher Scientific, Waltham, Massachusetts, USA). Grids were blotted for 5 s with a force of 5 (a setting on the Vitrobot) before being plunged into liquid ethane. For Muddy phage, three microliters of concentrated phage particles were added to a freshly glow-discharged Quantifoil R2/1 grid (Quantifoil Micro Tools GmbH, Großlöbicha, Germany) and plunge-frozen with a Vitrobot Mk IV into a 50:50 mixture of liquid ethane:propane^52^

### Cryo-Electron Microscopy

Data were collected on a 300 keV Titan Krios (Thermo Fisher Scientific, Waltham, Massachusetts, USA) at the Pacific Northwest Center for Cryo-EM with either a K3 or Falcon 3 direct electron detector (Gatan, Pleasanton, CA, USA). The data for Muddy was collected on a 300 keV Titan Krios 3Gi at the University of Pittsburgh with a Falcon 3 direct electron detector (Thermo Fisher Scientific, Waltham, Massachusetts, USA). Supporting Table 2 provides the collection parameters for each phage.

### Cryo-Electron Microscopy Data Analysis

Relion 3.1.1^53^ was used for phage capsid reconstructions using the standard workflow. CTF Refinement was performed using the default settings. Bayesian polishing was not performed since it made little improvement on resolution (approx. 0.1 Å for Bobi when attempted) for the computational time. Ewald sphere correction was carried out for each particle using the relion_image_handler command that is included with Relion. The mask_diameter value used in the Ewald sphere correction is reported in Supporting Table 2.

### De novo model building

The amino acid sequences of the major capsid proteins were folded with Alphafold^54^ version 2.0 using the default settings on a local workstation. The highest ranked prediction model was fitted into the cryo-EM map using ChimeraX^55^ version 1.3 and the “Fit in Map” command. Coot^56^ version 0.9.2 was then used to manually fit the model into the density using the “Stepped sphere refine active chain” provided by the python script developed by Oliver Clarke^57^. Any remaining protein backbone that was incorrectly placed was then manually moved into the correct density. All maps were of sufficient quality for side chains to be easily recognizable. The real-space refinement tool of Phenix^58^ version 1.19.2-4158 was used with default settings to refine the model and Coot was then used to manually fix the majority of the issues identified through Phenix. The final step was to use the ChimeraX plugin, Isolde^59^ version 1.3, to refine the major capsid protein model. The whole model simulation was used with a temperature of 20 °K. All other parameters were default. After the first model was completed, the asymmetric unit of the capsid was created using a similar workflow with a final Isolde refinement of the entire asymmetric unit.

### Phylogenetic analysis of the major capsid protein amino acid sequences

Amino acid sequences of the three major capsid protein phams (named as of July 2021: 4631, 15199, 57445) were downloaded from PhagesDB^60^ and merged into a single multifasta file. The divergent nature of these protein sequences required an alignment algorithm that could permit a large number of gaps in our multiple sequence alignment. To that end, we aligned the major capsid proteins using MAFFT (v7.453)^61^ with the following parameters: globalpair, unalignlevel 0.8, leavegappyregion, and maxiterate 1000.

A maximum likelihood phylogeny was created from the multiple sequence alignment using IQTree (v1.6.6)^62^ with the following parameters: ModelFinder Plus^63^ (-m MFP), and 100 non-parametric bootstraps (-bb 100). The model finder chose an LG model with empirical frequencies and five rate categories (LG+F+R5) as the most likely model based on the Bayesian information criterion. The resulting phylogeny was visualized in Figtree (v1.4.4)^64^. Nodes were collapsed only when the collapsed node contained a single pham from a single phage subcluster.

### Alphafold of major capsid proteins

To create the structural dendrogram, we used Alphafold to predict the three-dimensional protein fold of a representative major capsid protein from each cluster (139 total clusters), as well as every Singleton (62) major capsid protein. All protein sequences were obtained from the actinobacteriophage database (PhagesDB)^60^ and Phamerator^65^ in July 2021. For the few clusters (A, BN, CZ, DN, and F) that have more than one major capsid protein phamily, we folded a representative of each major capsid protein phamily from that cluster. There are forty-two annotated major capsid protein phamilies in the actinobacteriophages, spread across the 201 clusters and Singletons. In total, five clusters (DK, DS, EK, EM, and FC) and eight Singletons have no annotated major capsid protein and were therefore excluded from this analysis. The 18 total excluded phages account for less than 0.5% of the total number of annotated actinobacteriophages, so their exclusion is unlikely to skew the results. Cluster BO, which contains two phages, was also excluded from this analysis since they do not use the HK97-fold in their major capsid protein and are part of the Tectiviridae family of viruses. In total, 201 major capsid proteins were predicted with the default Alphafold settings and the major capsid protein amino acid sequence as input. The model with the highest confidence was used in the structural map. PDB files were manually truncated to remove the N-arm and the delta domain if present. The N-arm was truncated to approximately where the N-arm crosses behind the spine helix of the major capsid protein. The fasta files and PDB files of the predicted full-length and truncated major capsid proteins can be found in Supporting Information.

### Creation of a structural dendrogram using Homologous Structure Finder

In this study, we applied automatic structure alignment and the structure-based classification method Homologous Structure Finder (HSF)^66^, which allows comprehensive comparisons of proteins, not only within a protein family (such as RNA-dependent RNA polymerase)^67^ but also between protein families and superfamilies, significantly extending the depth of sequence-based phylogenies^66^. HSF identifies the equivalent residues for a pair of protein structures by comparing a set of amino acid properties (e.g., physiochemical properties of amino acids, local geometry, backbone direction, local alignment, and Cα distances)^66^. The two protein structures that are the most similar based on the properties are merged into a common structural core which then represents the pair in the later iterations. Next, the structure or a core from a previous iteration, best matching to an existing core or to a single structure not in any core yet, is merged either to a core or to another structure. The iterations are continued until all the protein structures are part of a clustering and a single structural core is identified for all the proteins in the data set. The equivalent residues in the structural core can be considered homologous, similar to high-scoring columns of multiple sequence alignment.

Pairwise comparison of the properties of the residues in the homologous positions of the common structural core between the original structures results in a pairwise distance matrix, which can be then used for constructing a structure-based distance tree^66^. The distances in such structure-based distance trees do not necessarily scale with respect to time, as changes in protein structure may not be continuous. However, the clustering of proteins in the structure-based distance tree constructed using HSF has been shown to follow the sequence-based classification of proteins into protein families, even when the common core contains less than 40 residues^67^. Thus, structure-based analysis is appropriate for a rough estimation of evolutionary events and relationships between protein families when the proteins share little or no detectable sequence similarity, and the accuracy of estimation of the evolutionary events increases as the sequence similarity increases.

### Data Deposition

PDB/EMDB/EMPIAR (unprocessed micrographs) accession numbers are as follows:

Adephagia: 8EC2/EMD-28012/EMPIAR-11200

Bobi: 8EC8/EMD-28015/EMPIAR-11201

Bridgette: 8ECI/EMD-28016/EMPIAR-11209

Cain: 8ECJ/EMD-28017/EMPIAR-11205

Che8: 8E16/EMD-27824/EMPIAR-11190

Cozz: 8ECK/EMD-28018/EMPIAR-11206

Muddy: 8EDU/EMD-28039/Not deposited in EMPIAR

Ogopogo: 8ECN/EMD-28020/EMPIAR-11207

Oxtober96: 8ECO/EMD-28021/EMPIAR-11208

Ziko: 8EB4/EMD-27992/EMPIAR-11195

## Results

### The actinobacteriophages have forty-two major capsid protein phamilies

There are currently (August 2022) over 4000 sequenced and annotated actinobacteriophages, which can be grouped into over 139 clusters and sub-clusters. Clustering is based on shared gene content between phage genomes, such that a phage is included in a cluster if it shares at least 35% of its genes with any member of that cluster (e.g. Cluster A, Cluster B, etc). Therefore, phages within a cluster are generally more globally similar to one another than to phages in other clusters. Some clusters can be similarly divided into sub-clusters (e.g. Subcluster A1, Subcluster A2, etc). Clusters range from having just two members (e.g. Cluster X) to over seven hundred (Cluster A), with most having fewer than fifty. Additionally, there are sixty-six “Singletons” (August 2022), those phages that have a genome that does not fit into an existing cluster. These cluster/subcluster/singleton groupings do not reflect firm biological distinctions, as phage genomes are pervasively architecturally mosaic, and phage populations likely span a continuum of diversity^68^.

The shared gene content comparison used for clustering is done at the protein level after genes have been translated and their products sorted into protein “phamilies” using Phamerator^65^ and a pipeline built on MMseqs2 (Gauthier and Hatfull, manuscript in preparation)^69^. A phamily^65^ is defined as a group of related proteins and although built with k-mer-based methods, proteins within a phamily typically have a minimum pairwise 20% amino acid identity. Amino acid sequence analysis of approx. 3200 major capsid proteins shows that there are forty-two major capsid protein families (termed phamilies/phams) within the actinobacteriophages database (July 2021). The majority of the 139 phage clusters have a single major capsid protein phamily and, as shown in this paper, it is possible to recapitulate most of the shared gene content-based phage clustering using only the major capsid protein.

### The F1 sub-cluster contains three major capsid protein phamilies

Because of the mosaic nature of phage genomes, some cluster/subcluster groups (e.g. A, BC, CZ, DN, and F) include multiple major capsid protein phamilies; Subcluster F1 has the most with three different major capsid protein phamilies. Previous structural studies with the *Escherichia coli* CUS-3 and *Salmonella* P22 phages have shown that major capsid proteins with minimal amino acid sequence identity (less than 15%) can result in almost identical capsid morphologies and HK97-folds^39, 46^. We, therefore, started the systematic investigation of the actinobacteriophages with the F1 major capsid proteins to address whether the three major capsid protein phamilies in the F1 subcluster are the same HK97-fold with highly diverged amino acid sequences, or whether they are three distinct folds.

In total, 180 phages within the F1 subcluster use one of three major capsid protein phamilies; the three major capsid protein phamilies show a minimum of 97% amino acid sequence identity within a phamily and a maximum of 15% between the phamilies.

Since the amino acid sequence identity is so high within each phamily, we chose a representative phage from each of the major capsid protein phamilies. These representatives are Che8 (phamily/pham number 4611, major capsid protein, gene number = 6), Bobi (phamily 15199, major capsid protein, gene number = 7), and Ogopogo (phamily 57445, major capsid protein, gene number = 8) for structural analysis. Note: phamily numbers are subject to change but are accurate at the time of writing (August 2022). Cryo-electron microscopy was used to determine a sub 3 Å map (Figure 2 A) for each of the three representative phages (see Supporting Table 2 for collection parameters, analysis, and final resolutions). The capsids all use the T=9 icosahedral architecture, with 540 copies of the major capsid protein, and are of similar size (740 Å diameter) and internal volumes (approx. 3x10^7^ Å^3^), which is expected since they package double-stranded DNA genomes of very similar length (Supporting Table 1). The high-resolution data obtained from our studies allowed for *de novo* model building of the major capsid protein’s amino acid sequence into the corresponding map. The resulting models of the three phages confirmed that each phage adopts the HK97- fold in their major capsid protein (Figure 2 B).

**Figure 2.**
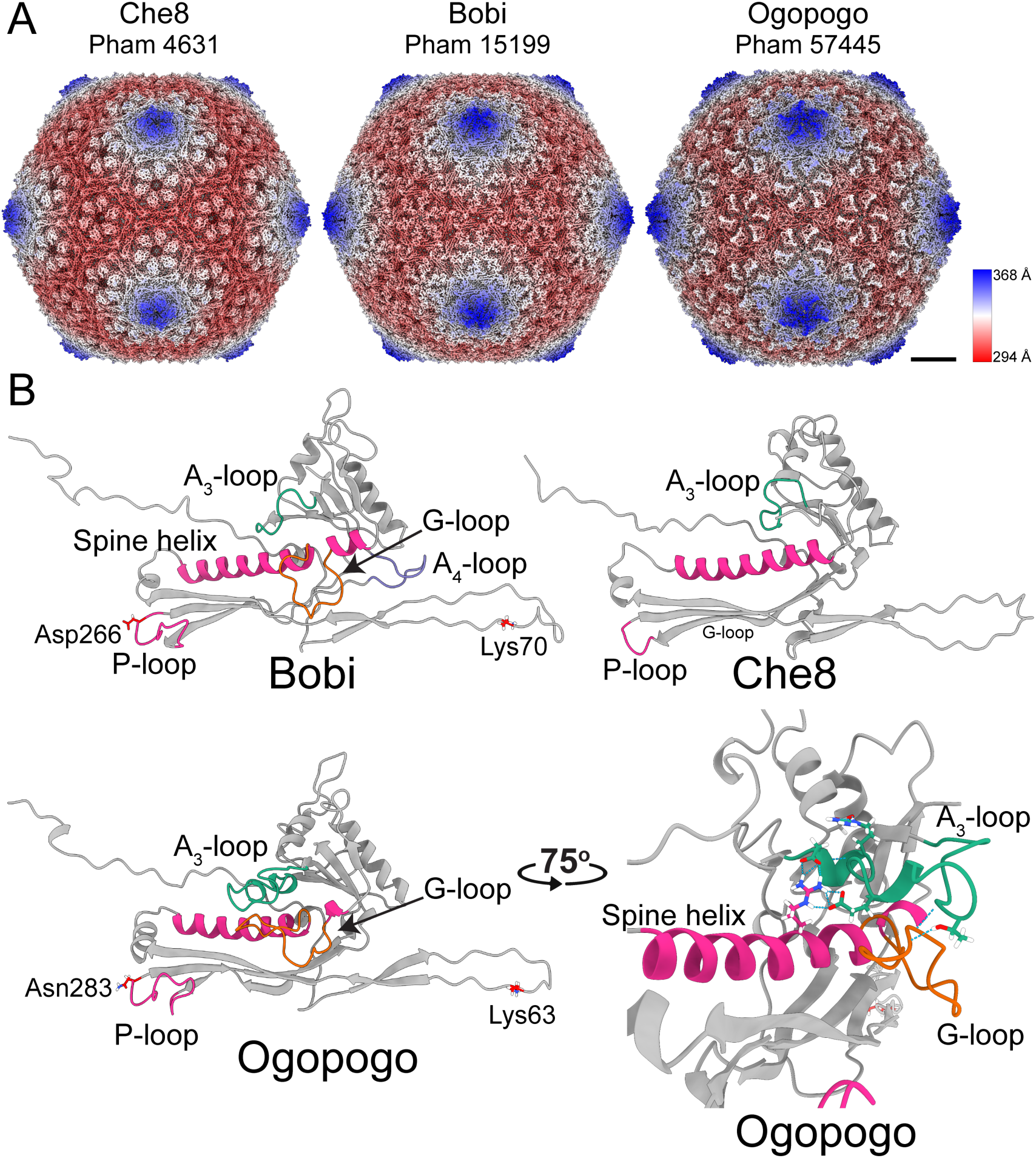
Structures of the F1 bacteriophages. A) Surface representations of cryo-EM maps of capsids from the three representative major capsid protein phamilies of the F1 subcluster: Che8, Bobi, and Ogopogo. Each phamily number is displayed below the bacteriophage name. The capsids have been colored by the radial distance from the center of the capsid (the color key is shown on the right-hand side). Scalebar = 10 nm. B) Models of the HK97-fold of each representative bacteriophage with select areas highlighted and labeled. The lysines and aspartic acid/asparagine involved in the isopeptide bond of Bobi and Ogopogo are also labeled (Che8 does not have an isopeptide bond). A zoomed-in and rotated image of the A_3_-loop and G-loop of Ogopogo is shown bottom right. Throughout the paper, the cryo-EM derived HK97- folds shown are of equivalent positions in the capsid and are the hexamer subunit adjacent to the pentamer subunit in the asymmetric unit.

A comparison of the three major capsid proteins, which have ∼15% amino acid sequence identity, revealed that the HK97-folds of the representative phages are very similar to one another (Figure 2), with Root Mean-Square Deviation (RMSD) of atomic position values <1.35 Å. While each fold is structurally similar to the original HK97-fold of the HK97 virus^23^, some key differences exist. Che8 lacks the G-loop that is found near the C-terminal end of the spine helix in the HK97 phage major capsid protein (Figure 1 A), therefore, Che8 has a continuous spine helix (Figure 2 B). Che8 also lacks the “protein chainmail” of the HK97 phage; the cryo-EM density map revealed that there was no density to suggest isopeptide bond formation, nor were there any amino acids in the correct location to potentially form an isopeptide bond. Unlike Che8, both Bobi and Ogopogo are more similar to HK97. They both contain a G-loop, although they are more extended when compared to the original HK97 fold. Ogopogo additionally has an extended A-loop that extends over the G-loop and makes important stabilizing contacts with the G-loop and spine helix (Figure 2 B, bottom right). The A-loop is in the same position as the A-loop of phage T7 where it was first characterized^26^; we name this the A_3_-loop. Bobi has an additional loop between the A and P domains, in a similar position to the A-pocket described in phage T7^26^; we name this the A_4_-loop. Ogopogo and Che8 do not have the A_4_-loop. The A_4_-loop in Bobi does not make intermolecular interactions with other adjacent major capsid proteins, although it does make some intramolecular hydrogen bonds. However, the A_4_-loop is spatially very close to the A_3_-loop of an adjacent major capsid protein (discussed in more detail later). Furthermore, both Bobi and Ogopogo have an isopeptide bond, with clear density in the cryo-EM map showing the covalent link between a lysine (Bobi, lysine 70 and Ogopogo lysine 63) in the E-loop and either aspartic acid or asparagine in the P-domain (Bobi, Asp 266 and Ogopogo Asn 283) of the adjacent major capsid protein: this demonstrates that they form the characteristic “protein chainmail” like the original HK97 fold. Ogopogo and Bobi both have an extended P-loop within the P-domain and three of these are found in close contact around the local 3-fold axis of the capsid. These P-loops will be discussed in greater detail later. This structural comparison of the three F1 major capsid protein phamilies suggested that the Che8-like phages may be more divergent from Bobi and Ogopogo.

For simplicity, from this point on the major capsid protein phams will be called by the corresponding representative phage used in Figure 2; the Che8-like phages (pham 4631); the Ogopogo-like phages (pham 57445) and the Bobi-like phages (pham 15199).

### The three major capsid protein phamilies of the F1 subcluster constitute two structural groups of major capsid protein in the actinobacteriophages

To confirm the structural observations, we next put the three F1 major capsid protein phamilies into the broader context of the major capsid proteins from the actinobacteriophages database since focusing on just the three F1 phamilies would likely not reveal much insight into their level of evolutionary relationship due to their relatively high structural similarity. We, therefore, created a structural dendrogram of all the major capsid protein phamilies annotated in the actinobacteriophages (Figure 3).

**Figure 3.**
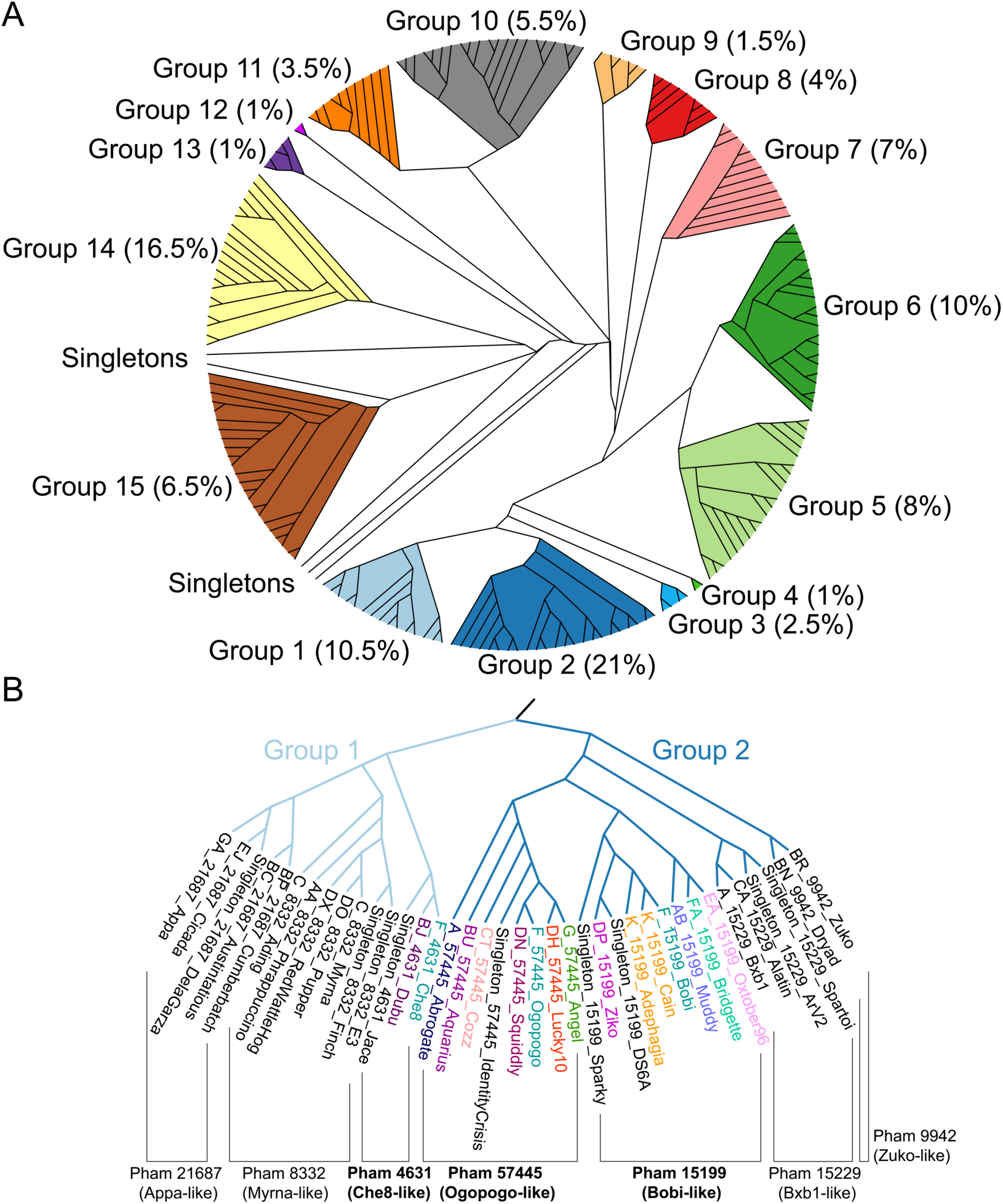
Structural dendrogram of the actinobacteriophages. A) 201 Alphafold- predicted representative major capsid protein HK97-folds, representing 99% of the actinobacteriophages, were clustered based upon their structural similarity using the Homologous Structure Finder algorithm. Each cluster is designated with a Group number, and the percentages in parenthesis after the group number are the number of actinobacteriophages (from the 4000+ in phagesdb.org) that are found within each structural group. The map is colored to highlight each structural group. B) A zoomed- in image of the structural dendrogram highlighting Structural Groups 1 and 2 that contain the three F1 major capsid protein phamilies. Representative bacteriophage names are shown and are colored by cluster. Within Groups 1 and 2 are subgroups of proteins from the same phamily, as indicated.

Previously, structural comparison of distantly related, yet conserved, protein folds has been used successfully to imply evolutionary links between viral capsid proteins; for example, with the PRD1 and other double jelly-roll viral capsid proteins including adenovirus^66, 70, 71^, as well as showing a link between the dsDNA tailed phages and Herpes virus^3^.

To create the structural dendrogram we used Alphafold^54^ to predict the three-dimensional HK97-folds of the major capsid proteins. Folding every major capsid protein in the actinobacteriophage database (over 3000 entries when the analysis was carried out in July 2021) was not feasible from a computational standpoint due to the large number of major capsid proteins. Therefore, we selected a representative major capsid protein from each cluster (139 total clusters at the time of analysis), as well as for every Singleton (62 at the time of analysis), as each Singleton could represent a future cluster distinct from the extant groups. For those clusters (A, BN, CZ, DN, and F) with more than one major capsid protein phamily, we folded a representative of each major capsid protein phamily from that cluster. In total, the structure of 201 major capsid proteins were predicted using Alphafold and represent the forty-two annotated major capsid protein phamilies of the actinobacteriophages.

We validated a subset of the Alphafold predictions with cryo-EM derived structures (Supporting Figure 1), revealing excellent agreement for most of the HK97-fold (RMSD values between 0.8 - 1 Ångstrom). While Alphafold performs well for most of the fold, including the structured A-domain and P-domain, it does not as accurately predict the E-loop, and in most cases, the N-arm is badly predicted (Supporting Figure 2). These results concerning the E loop and N arm are understandable: the N-arm and E-loop make long-range interactions with other major capsid proteins within the capsid and previous studies have shown quite different conformations of the same HK97-fold major capsid protein within its asymmetric unit and during its assembly, showing that both the E-loop and the N-arm are often in different positions and, consequently, hard to predict.

All tailed phages use a scaffolding protein domain during capsid assembly to create the empty capsid into which the viral DNA genome is packaged^72^. This scaffolding protein domain can be either an independent protein (called scaffolding protein) or an N-terminal extension of the major capsid protein sequence (called the delta domain) and in phage HK97 is cleaved from the major capsid protein after capsid assembly^72^.

Approximately 35% of the Alphafold predicted major capsid proteins from the actinobacteriophages had an N-terminal extension that was similar in size to the HK97 phage delta domain. Some of the predicted delta domains in the actinobacteriophage major capsid proteins were almost as large (300 amino acids) as the major capsid protein and it is not possible to predict the cleavage site between the delta domain and the post-cleavage N-arm. We therefore manually truncated all the Alphafold predicted major capsid proteins to remove the N-terminal arm and the delta domain, if present, to make sure that we did not introduce bias from the N-arm and delta domains into the structural comparison. PDB-format files for the 201 full-length and truncated major capsid protein predictions can be found in Supporting Information.

These 201 truncated major capsid protein predictions were then used to create a major capsid protein structural dendrogram of the actinobacteriophages using the Homologous Structure Finder algorithm^66^ (Figure 3 A). The algorithm compares the three-dimensional Alphafold predictions and classifies the major capsid proteins on their structural similarity. The sophisticated classification methodology allows for the creation of a structural dendrogram whereby common structural elements between the major capsid proteins are identified and a common structural ancestor can be inferred. It has been used successfully for other protein fold lineages^66, 73^. The major advantage of this methodology is for detecting similarities in protein folds even when no similarities remain in the amino acid sequences. The forty-two major capsid protein phamilies within the actinobacteriophages have less than 20% amino acid sequence identity, meaning that comparison of the major capsid protein structures is the best way to reveal similarity and infer evolutionary links between the major capsid proteins.

The structural map of the actinobacteriophages revealed that the 42 major capsid protein phamilies can be classified into 15 structural groups (Figure 3 A) that are likely to be evolutionarily related. Each structural group is classified solely on the three-dimensional structure of the major capsid protein and is independent of the cluster and major capsid protein phamily. Beyond the 15 structural groups are several “structural Singletons”. The structural dendrogram supports the initial structural comparison of the three Cluster F1 phage major capsid proteins in that the Che8-like phamily, sorted into Group 1, is more diverged from the Ogopogo and Bobi-like phamilies found in Group 2 (Figure 3 B). Both groups are relatively large and just over 30% of all the annotated actinobacteriophages use major capsid proteins found in these structural groups (Group 1: 10.5% of all annotated actinobacteriophages and Group 2: 21%).

Group 1 (Supporting Figure 3) contains three different major capsid protein phamilies: 4631 (Che8-like), 8332 (Myrna-like), and 21687 (Appa-like). It is very similar to the HK97-fold of the HK97 virus, with RMSDs when compared to the HK97 major capsid protein of between 1 and 1.3 Å. However, all of the Group 1 major capsid proteins lack the G-loop of the spine helix. The Appa-like (21687) sub-group all have a 7-strand beta-sheet in the A domain, compared to the 5-strand sheet of the HK97-virus. The Appa-like members also have a large loop of varying length at the very end of the spine helix. The Myrna-like (pham 8332) sub-group all have a 6-strand beta sheet in the A domain. The Che8-like (pham 4631) sub-group all have the canonical five-strand beta sheet in the A domain but have an extra helix at the top of the A domain and an elongated spine helix with two extra turns when compared to the HK97 spine. All three structural sub-groups are defined by their major capsid protein phamily with no evidence of a member of a phamily being more closely related to another phamily than its own phamily. For example, no Che8-like major capsid proteins cluster within the Appa-like or Myrna-like structural sub-groups.

Group 2 (Supporting Figure 4) contains four different major capsid protein phamilies: 9942 (Zuko-like), 15229 (Bxb1-like), 15199 (Bobi-like), and 57445 (Ogopogo-like). The main architecture of the HK97-fold of Group 2 is once again highly similar to the HK97- fold of phage HK97, with an RMSD of between 1 and 1.4 Å. The main difference between Group 2 and HK97 is the presence of a larger G-loop within Group 2. The Ogopogo-like (57445) phages all have an elongated A_3_-loop within the A-domain directly above the G-loop and the A_3_-loop is predicted to interact with the G-loop via hydrogen bonding; this is confirmed by the cryo-EM-derived model of Ogopogo. The Bxb1-like (15229 sub-group) and Bobi-like (151999 sub-group) proteins contain an elongated P-loop in one of the beta-sheets of the P-domain. This loop is typically located at the 3-fold axis of the capsid and suggests that these phages have a raised “turret” at the three-fold. The “turret” can be seen in the cryo-EM maps of the Bobi-like (15229) capsids. The final sub-group, the Zuko-like (9942), are the most distantly related to the other three, although, like the Ogopogo-like sub-group, they have a similar extended A_3_-loop.

**Figure 4.**
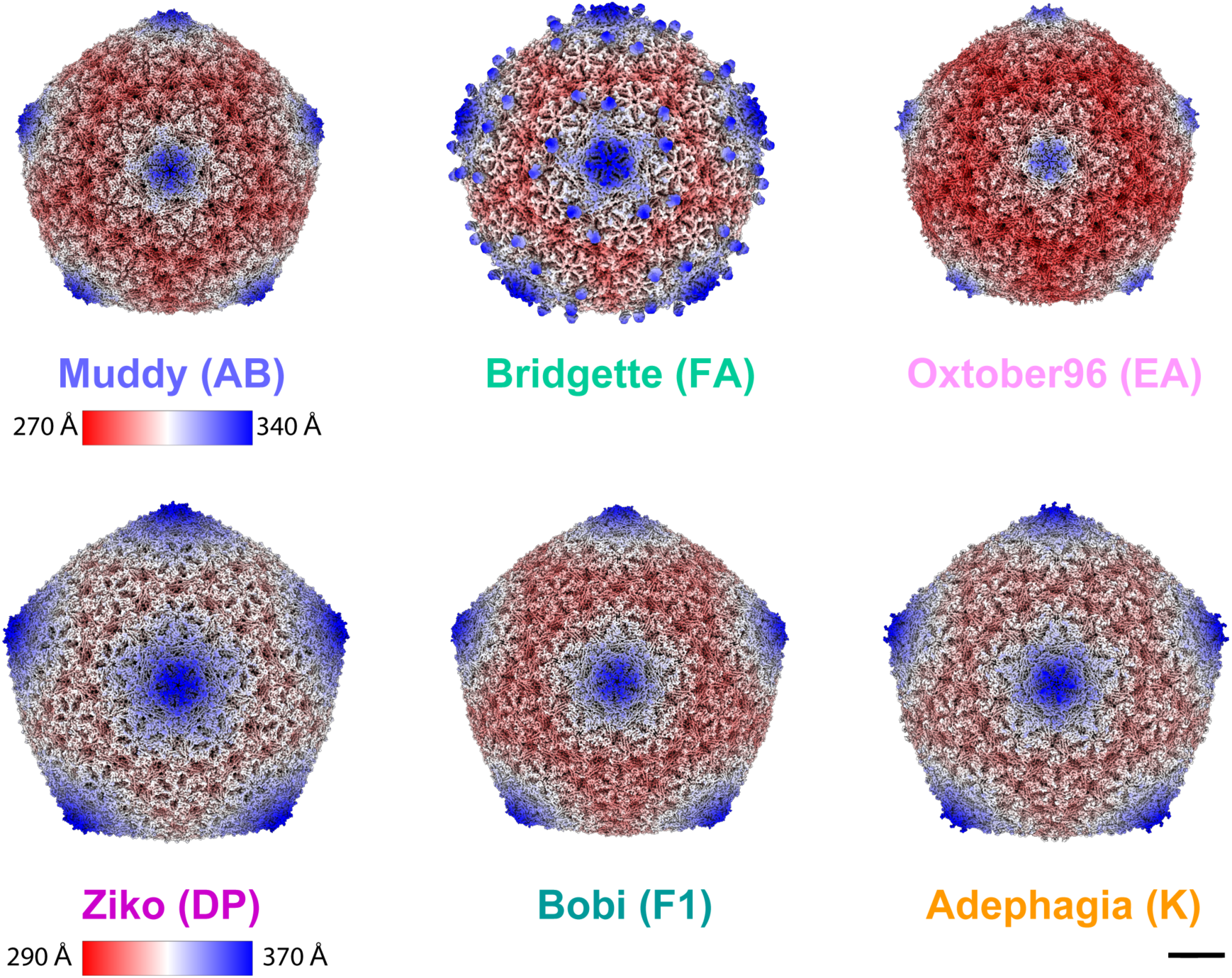
Cryo-EM maps of representative bacteriophages from the Bobi-like (15199 phamily) of major capsid proteins. Muddy, Bridgette, and Oxtober96 all form T=7 capsids. Ziko, Bobi, and Adephagia all form T=9 capsids. Bridgette has a decoration protein that assembles as dimers around the pentamer. The scale bar is 10 nm. Maps are radially colored. Muddy, Bridgette, and Oxtober use one radial color scheme, and Ziko, Bobi, and Adephagia another. For clarity, names are colored by cluster using the same colors as in Figure 3. The bacteriophage cluster name is shown in brackets after the bacteriophage name. The Bobi map is reproduced here from Figure 2 for comparison with the other five cryo-EM maps. Scalebar = 100 Å.

### The Bobi-like (15199) phamily can form differently-sized capsids

Having characterized the three F1 phamilies and shown that Che8 is likely to be evolutionarily distinct from the other two phamilies, we next wanted to investigate how the HK97-fold and capsid stability mechanisms are conserved within closely related phages. We chose to concentrate on the Group 2phages since they use the protein “chainmail” found in HK97. Alignment of all the Alphafold-predicted major capsid proteins from Group 2 shows lysine and aspartic acid/asparagine at the expected positions in the E-loop and P-domain in the majority of the Group 2 phages, apart from those in the Zuko-like (9942) sub-group and a subset of the Cluster K phages in the Bobi-like (15199) sub-group. Likewise, all the sub-groups, apart from the Zuko-like (9942) have a glutamic acid in a position within the P-domain that could act like the Glu363 in the HK97 major capsid protein for the catalysis of the cross-linking.

It was surprising to identify some of the K subcluster lacking the lysine residue needed for the isopeptide bond since all the other Bobi-like (15199) phages were predicted to use it. Removal of the isopeptide bond in HK97 by mutating the E-loop lysine to tyrosine results in non-viable phage particles indicating that the isopeptide bond is required^74^ for infectious capsids to be made. The Bobi-like (15199) phages, therefore, provided an opportunity to characterize phage structures with and without the isopeptide bond and investigate how/and if the capsid compensates for its absence. We carried out cryo-EM on five other members of the Bobi-like (15199) phage capsids to yield density maps of < 4 Å resolution (Supporting Table 2). We found that some of the Bobi-like (15199) phages formed T=9 capsids while others formed T=7 capsids (Figure 4). In each of the six phages (including Bobi), only the major capsid protein was identified in the cryo-EM map; no minor capsid proteins were observed. Bridgette of Cluster FA, however, did have a decoration protein (Gp7, Supporting Figure 6) that bound as dimers around the 5-fold axis of the pentamer, reminiscent of the phi29 phage spike proteins^28^.

**Figure 6.**
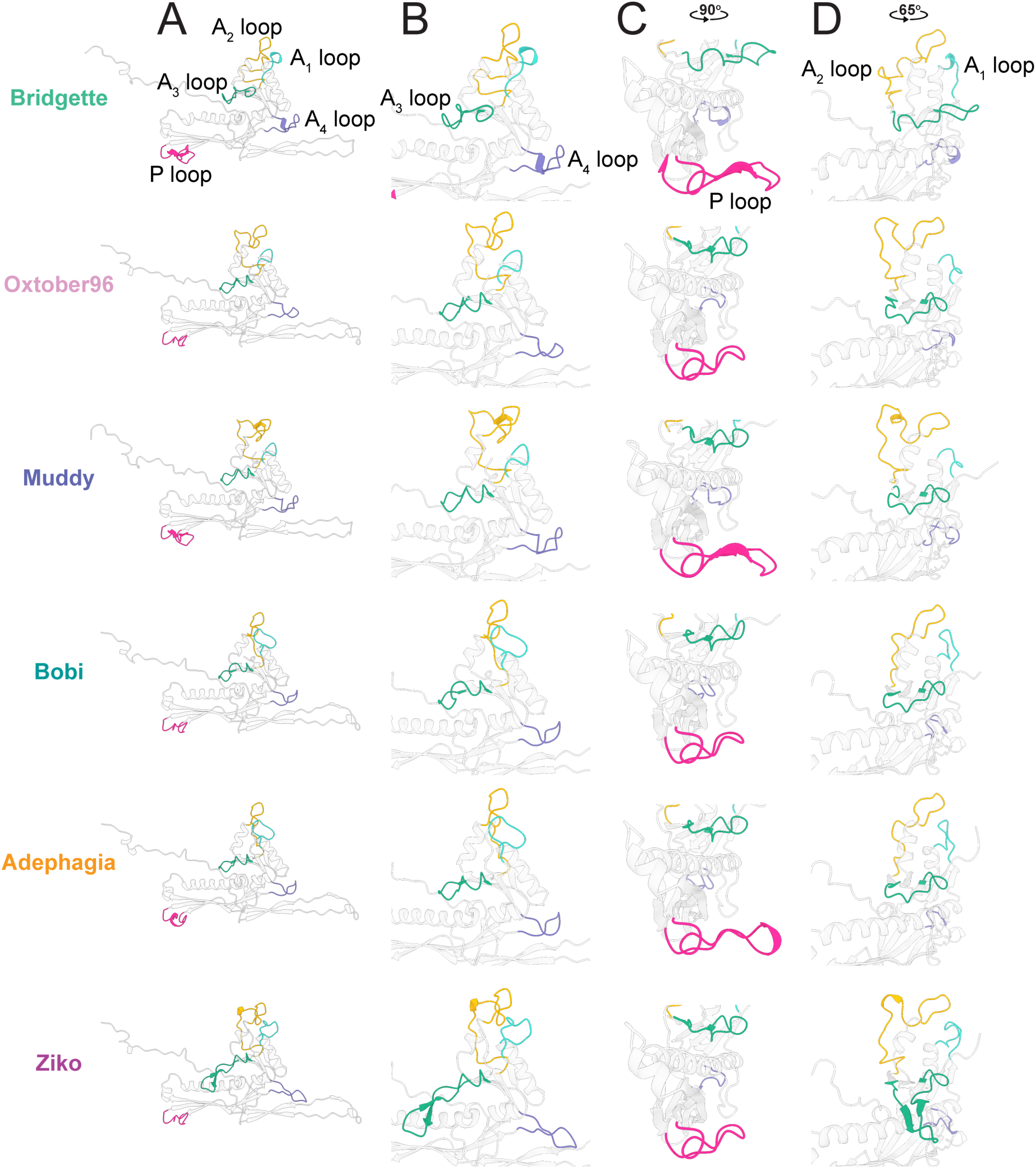
The variable loops of the Bobi-like (15199) major capsid proteins. A) Major capsid protein subunits for each phage with all the variable loops highlighted. B, C, and D show zoomed-in images of the same major capsid protein as in A.

### The HK97-folds of the Bobi-like phamily (Pham 15199) are highly conserved, but exhibit structural diversity in the loop regions

The high resolution of the six maps of the representative Bobi-like (pham 15199) phages allowed for *de novo* model building of each of the major capsid proteins. Comparison of the HK97-folds showed very high similarity between them (Figure 5), with the highest RMSD value of 1.2 Å (Supporting Table 3). The protein fold is highly conserved and near identical, especially in the secondary structure alpha helices and beta sheets and can be overlaid without much deviation along the protein fold. It is within five loop regions that the most structural diversity is observed (Figure 6). We have designated these loop regions A_1_-A_4_ because of their position in the A domain, as well as the P-loop found in the P-domain. The A_3_ and A_4_ loops were first described in the HK97-fold of the phage T7 major capsid protein and named the A-loop (A_3_ loop) and A-pocket (A_4_ loop)^26^. We have renamed them here because of the extra loops we have identified. All the structurally characterized Bobi-like (pham 15199) phages have both A_3_ and A_4_ loops. Within Bobi, Oxtober96, and Adephagia, the two loops are of similar length and make intramolecular interactions but do not interact with one another. The other three phages, Bridgette, Muddy, and Ziko, all have increasingly long A_3_ and A_4_ loops, with those of Ziko being the longest (Figure 6 B). The extended A_3_ loop in Ziko extends out over its G-loop, reminiscent of the same loop found in the Ogopogo-like phages (Figure 2 B). The G-loop stabilizes the Ziko A_3_ loop with a salt bridge involving Gln130 in the G-loop and Asp223 in the A_3_ loop. This is not observed in the Ogopogo- like phages where hydrogen bonds are used to stabilize the A_3_ loop above the G-loop (Figure 2 B). The extended A_3_ and A_4_-loops in Bridgette, Muddy, and Ziko stabilize the capsid through intermolecular interactions (Figure 7 A), either forming a single salt bridge (in the case of Bridgette) or a salt bridge combined with hydrogen bonds (in the case of Ziko) between the two loops on neighboring chains. Muddy does not make an intermolecular contact between the A_3_ and A_4_-loops, instead, the A_4_ loop forms a salt bridge with the N-arm of the adjacent major capsid protein. Only a single tryptophan (W224 in Bobi) is conserved in the A_3_ loop across the Bobi-like (15199) major capsid proteins. It forms a pi-pi interaction with a conserved phenylalanine (Phe127 in Bobi) that presumably stabilizes the A_3_ loop (Supporting Figure 6). No residues are conserved in the A_4_ loop.

**Figure 5.**
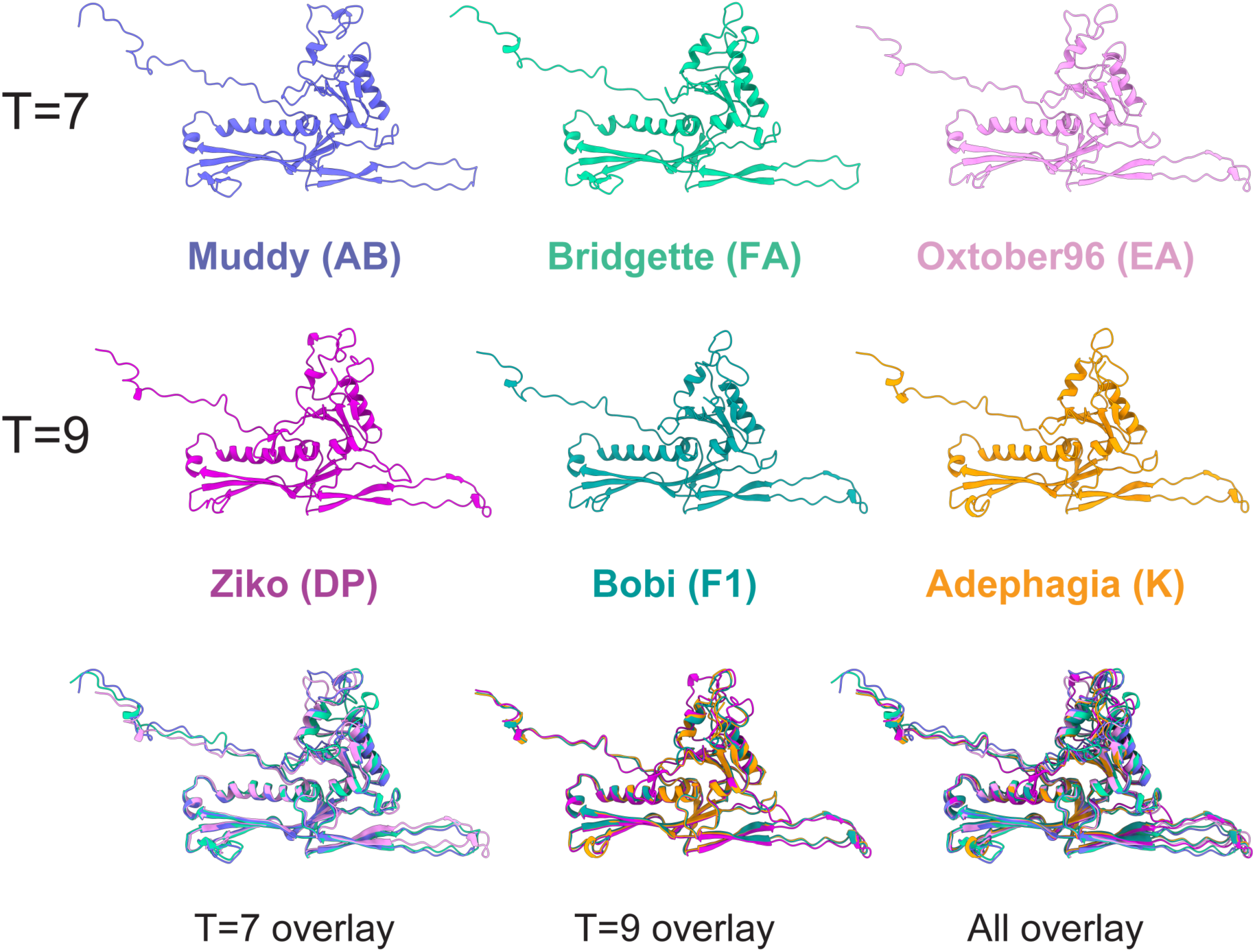
Models of the representative bacteriophages from the Bobi-like (15199) phamily of major capsid proteins. Model of each major capsid protein derived from the cryo-EM maps. Each model is of the hexamer subunit adjacent to the pentamer subunit, so models are directly comparable. Color coding is by phage cluster and is consistent with previous figures. The T=7 forming capsids (Muddy, Bridgette, and Oxtober96) are overlaid, as are the T=9 forming capsids (Ziko, Bobi, and Adephagia). All six models are also overlaid (All overlay).

**Figure 7.**
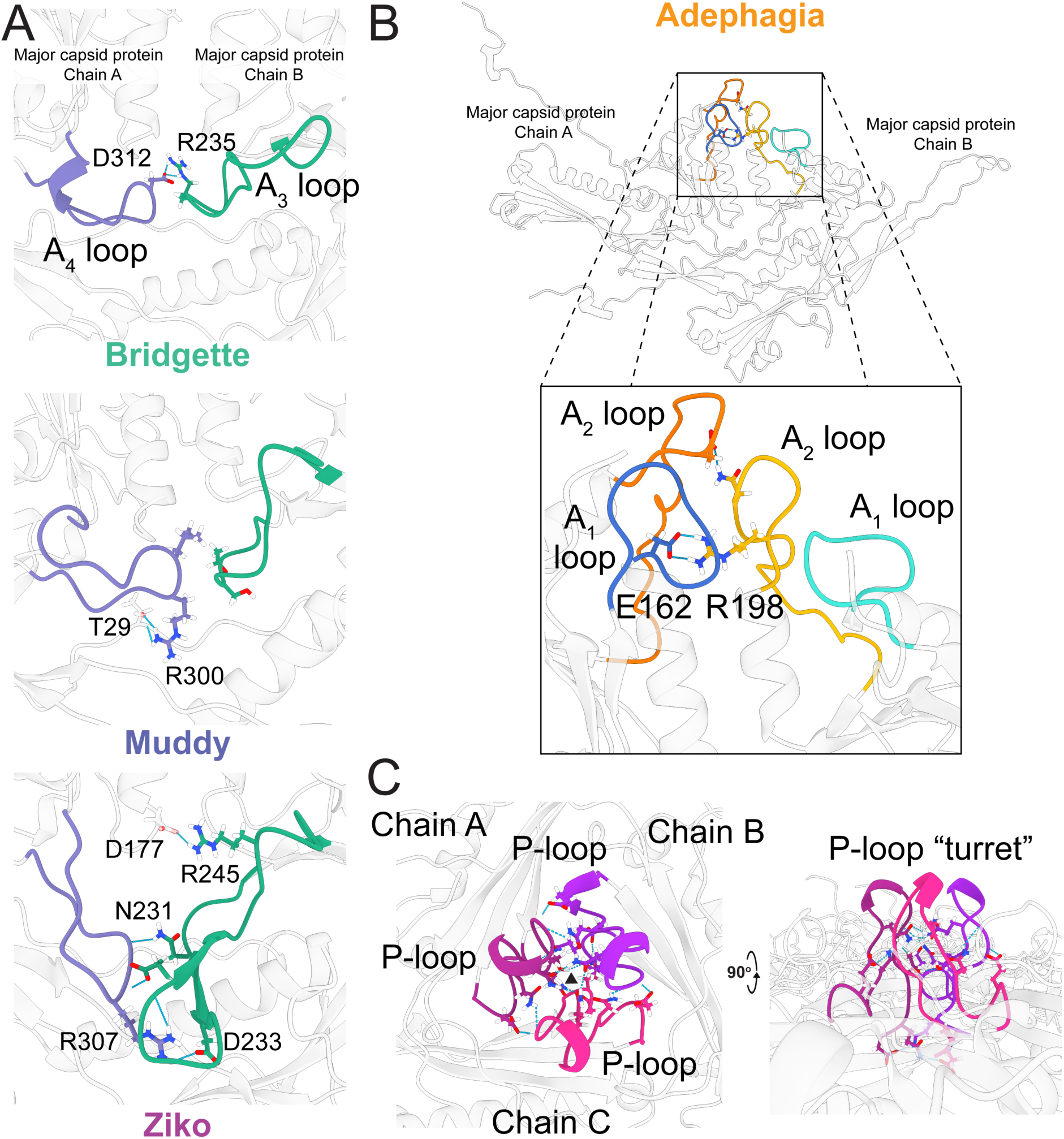
Intermolecular interactions between Bobi-like (15199) major capsid proteins. A) Interactions between the A_3_ and A_4_ loops of one major capsid protein subunit with an adjacent subunit for phages Bridgette, Muddy, and Ziko. B) The interaction between the A_1_ and A_2_ loops of one major capsid protein subunit and an adjacent major capsid protein in Adephagia (top) and a zoomed-in viewpoint of the same structure with the intermolecular interactions between sidechains shown and the conserved salt bridge highlighted with a black box (bottom). The Arginine/Glutamic acid salt bridge between the A_2_ loop (R198) and the A_1_ loop (E162) is shown. C) The P-loops of three major capsid protein subunits in Adephagia are colored in three different shades of magenta. The black triangle shows the center of the local 3-fold axis.

Extending this analysis to the wider Structural Group 2 (Figure 3) using the Alphafold predicted models (Supporting Figure 4) suggests that the two Singletons, DS6A and Sparky, both have elongated A_3_ and A_4_ loops like Ziko but the A_3_ loop forms a beta hairpin structure. The A_4_ loop is not universal amongst the Group 2 phages (Supporting Figure 4). For example, the Ogopogo-like (57445 phamily) and Bxb1-like (15229 phamily) phages, completely lack the A_4_ loop although they still have the A_3_ loop.

The A_1_ and A_2_ loops are found at the top of the A domain (Figure 6 D), with the A_2_ loop inserting into the center of the hexamer or pentamer capsomere. Oxtober96, Ziko, and Muddy all have elongated A_2_ loops. A comparison of the A_1_ and A_2_ loops in the context of the capsid (Supporting Figure 7) shows that there is a conserved salt bridge between an Arginine in the A_2_ loop of one major capsid protein and a Glutamic acid in the A_1_ loop of an adjacent major capsid protein (Figure 7 B). However, in Bridgette, the salt bridge is between the two A_2_ loops (Supporting Figure 7). The elongated A_2_ loops do not appear to result in a consistent increase in intermolecular or intramolecular interactions between the major capsid proteins (Table 2) and no other amino acids are conserved.

**Table 2.** Hydrogen bonds between the A_1_ and A_2_ loops. Hydrogen bonds and salt bridges specific to the A_1_ and A_2_ loops were determined in ChimeraX^49^.

| *Bacteriophage name* | *Intermolecular bonds* | *Intramolecular bonds* |
| --- | --- | --- |
| *Adephagia* | 3 | 32 |
| *Bobi* | 5 | 32 |
| *Ziko* | 6 | 31 |
| *Muddy* | 8 | 24 |
| *Oxtober96* | 8 | 25 |
| *Bridgette* | 4 | 43 |

The P-loop (Figure 6 C) makes contact with other P loops around the 3-fold axis, creating small “turrets” that stick outward from the capsid (Figure 7 C). The P-loops make several hydrogen bonds and salt bridges (Table 3) between adjacent P-loops that are likely to play a role in the stabilization of the local 3-fold axis. Adephagia has one of the longest P-loops and makes the most salt bridges and hydrogen bonds between the P loops out of all six Bobi-like phages.

**Table 3.** Hydrogen bonds in the local 3-fold axis. Hydrogen bonds and salt bridges specific to the P-loop in a single local 3-fold axis were assessed in ChimeraX^49^. The number of bonds between all the major capsid proteins that are involved in the local 3- fold was determined using PDBsum^64^.

| *Bacteriophage name* | *P-loop hydrogen bonds* | *P-loop salt bridges* | *3-fold hydrogen bonds* | *3-fold salt bridges* |
| --- | --- | --- | --- | --- |
| *Adephagia* | 41 | 9 | 207 | 73 |
| *Bobi* | 18 | 3 | 201 | 90 |
| *Ziko* | 18 | 3 | 236 | 89 |
| *Muddy* | 24 | 0 | 273 | 65 |
| *Oxtober96* | 16 | 0 | 246 | 108 |
| *Bridgette* | 19 | 3 | 198 | 79 |

The G-loop structure is well conserved across all of the Bobi-like (15199) phages and is a clear structural marker for this phamily of phage major capsid proteins despite only having a single conserved glycine and proline across the Bobi-like (15199) major capsid proteins (Supporting Figure 4).

### Covalent crosslink residues are not found in all of the Bobi-like major capsid proteins

Alignment of the Bobi-like (15199) phamily major capsid protein amino acid sequences revealed that a subset of the Cluster K phages lack the lysine in the E-loop at position 70 of the Bobi major capsid protein. This lysine, present in every other Bobi-like phamily member, as well as most of the Cluster K phages, is half of the amino acid pair that creates the isopeptide bond between two major capsid protein subunits. This lysine is substituted by isoleucine at position 70 (K70I, Figure 9 A) in 48% (78 in total) of the Cluster K phages, with the remaining Cluster K phages having the lysine at the correct position, demonstrating that it is not a rare occurrence. MAFFT alignment and phylogenetic analysis of the major capsid proteins showed that the Cluster K phages (Supporting Figure 8) are most closely related to the Cluster F phages (of which Bobi is the representative) with 70% amino acid sequence identity between Bobi and Adephagia (Clusters F and K, respectively). Within the K70I sub-group of Cluster K phages, there are nearby lysine residues that we hypothesize could act as the lysine for the isopeptide bond. Depending on the position of this misplaced lysine, these Cluster K phages can be divided into two groups (Figure 8 A). We have termed one group the Adephagia-like phages, and the other the Cain-like phages. Both contain the isoleucine replacement of the isopeptide-forming lysine. However, the Adephagia-like phages have a lysine adjacent to the isoleucine, while the Cain-like phages have a lysine four residues downstream (towards the C-terminus) from the isoleucine. The Cain-like phages are a relatively small group, with only eight members (∼5% of the Cluster K phages). Overall, the major capsid proteins of Adephagia and Cain are highly similar to the other Cluster K phages with an amino acid sequence identity of 92%. Having already obtained a high-resolution map of Adephagia (Figure 5), we performed cryo-EM on Cain and obtained a sub 3 Å map (Supporting Figure 9 A) to investigate whether it could form the isopeptide bond with the downstream lysine.

**Figure 8.**
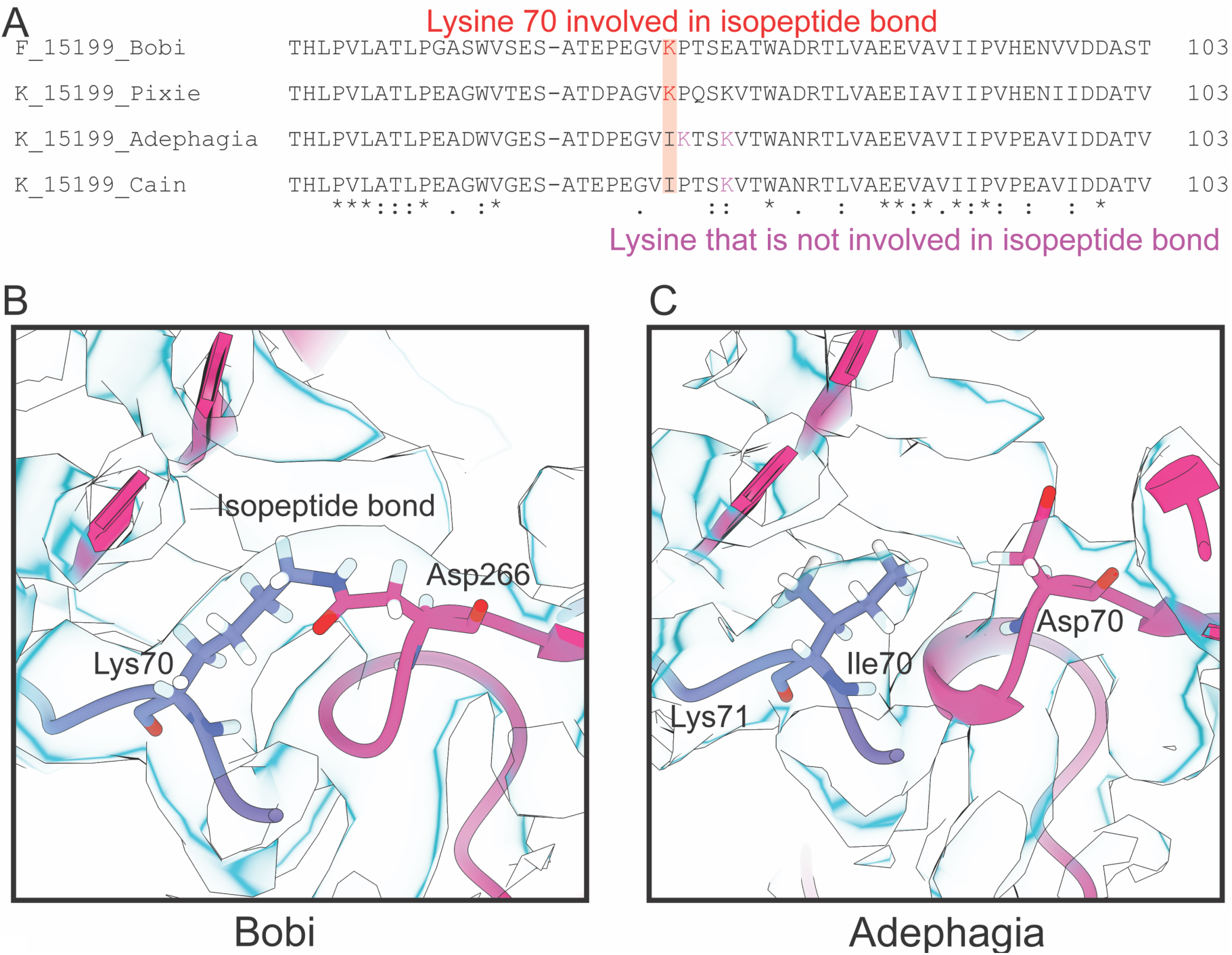
The isopeptide bond. A) Alignment of Bobi and the three main groups of the K cluster. B) Bobi isopeptide bond is shown with the cryo-EM map shown in light blue. C) Adephagia is shown that lacks the isopeptide bond.

Analysis of the six cryo-EM maps from the Bobi-like (pham 15199) phages (Figure 4), as well as the map of the Cluster K Cain phage confirmed that all the phages, except Adephagia and Cain, had clear density for the isopeptide bond between the lysine in the E-loop and an aspartic acid in the P-domain of the adjacent major capsid protein (Figure 8 B). The lysine residues within Adephagia and Cain that we hypothesized could form the isopeptide bond were therefore found not to be involved in covalent bond formation (Figure 8 C and Supporting Figure 9 B).

A consequence of the isopeptide bond is that all the major capsid proteins are covalently linked to one another, forming a large complexes. Previous SDS-PAGE analysis of the capsid of the HK97, where the isopeptide bond was first demonstrated (Figure 1), showed that the major capsid protein was unable to enter the gel due to its extensive crosslinking and large size^75^. We, therefore, ran the Ziko (that contains the isopeptide bond) and Adephagia (with no isopeptide bond) capsids on SDS-PAGE (Supporting Figure 10). The gel confirmed the cryo-EM analysis of isopeptide bond formation: no major capsid protein band was observed for Ziko at the expected molecular weight of 34 kDa but instead, there were dark areas at the top of the gel reminiscent of the HK97 SDS-PAGE analysis. Conversely, a large band consistent with the major capsid protein of Adephagia was detected on the gel at approximately 32 kDa (the predicted size of Adephagia gp13 major capsid protein is 32.7 kDa).

### The isopeptide bond catalytic site in the Bobi-like phages is similar to that found in HK97

Phage HK97 was the first (and until now, only) structurally characterized example of the isopeptide bond in the tailed phages; the bond is formed between lysine 169 and asparagine 356 (Figure 1) and has been well characterized. The isopeptide bond has also been shown biochemically in other phages, for example, D3^76^, Q54^77^, and L5^78^ but these have not been structurally characterized.

The formation of the isopeptide bond is catalyzed by a glutamic acid (Glu363 in HK97) located within a hydrophobic pocket made up of three amino acids (Val163, Met339, and Leu361)^24^ (Figure 1). The Bobi-like isopeptide bond (Figure 8 B) is similar to that found in HK97; it is formed between lysine 70 in the E-loop and aspartic acid 266 in an adjacent major capsid protein subunit. A highly conserved glutamic acid (Glu86) and valine (Val59) are found in almost identical structural positions as those in HK97.

However, there are no equivalent hydrophobic residues in Bobi that would recapitulate the hydrophobic pocket provided by HK97 Met339 and Leu361, suggesting that these residues are not conserved across all phages that use the isopeptide bond and that different ways to create the hydrophobic pocket are possible. All the identified amino acids within the Bobi-like phage catalytic site are highly conserved across all of 15199 pham apart from Lysine 70, which is substituted only in the aforementioned subset of the Cluster K phages: the Adephagia-like and Cain-like.

The catalytic glutamic acid is not structurally conserved across Group 2. In the Bobi-like (15199) sub-group the glutamic acid is structurally conserved (relative to HK97). In the same position within the other members of Group 2, all have a lysine at the same position. The Zuko-like (pham 9942) have no glutamic acid nearby that could play a catalytic role. The remaining members of Group 2 (Ogpogo-like and Bxb1-like) all have a structurally conserved glutamic acid nearby that could play a similar catalytic role. The position is confirmed in the cryo-EM map of Ogopogo with Glu290 in a position to possibly be the catalytic glutamic acid.

### Structural groups do not have a conserved hydrogen bond network

The lack of an isopeptide bond in Adephagia and the other Cluster K phages raised the question as to how they compensate for the absence of the covalent isopeptide bond around the local 3-fold axis for capsid stability. Removal of the isopeptide bond in HK97 results in unviable capsids, suggesting it is critical for the survival of virion. With no minor capsid protein or other accessory protein to compensate for the loss of the isopeptide bond, we examined the hydrogen bonds and salt bridges around the local 3- fold axis, hypothesizing that there would be an increased number of such interactions around the local 3-fold of Adephagia when compared to Bobi, its closest relative in the Bobi-like (15199) pham. Specifically, we predicted that the P-loops at the center of the 3-fold would have increased interactions in Adephagia. Indeed, this is what we observed, with an increase in both salt bridges and hydrogen bonds between the P-loops, with Adephagia having three times the salt bridges and double the number of hydrogen bonds when compared to Bobi (Table 3). This equates to an increase of 240 kJ/mol free energy between Adephagia and Bobi at the site of the P-loops, although this must be contrasted with a loss of 1068 kJ/mol in free energy because of the removal of the isopeptide bonds. However, despite the increase in interactions at the P-loop, which is located at the center of the local 3-fold axis, there was no increase in the number of hydrogen bonds and salt bridges between Adephagia and Bobi around the wider local 3-fold axis that takes into account all nine interacting major capsid proteins (Table 3 and Supporting Figure 11). We next expanded the analysis to the other members of the Bobi-like (15199) pham, which revealed that all the capsids had similar numbers of hydrogen bonds and salt bridges around the local 3-fold axis but only a handful of these were structurally conserved, with different distribution patterns of the interactions for each phage (Table 3 and Supporting Figure 11). We, therefore, conclude that extra stabilization around the local 3-fold axis is not required to compensate for the missing isopeptide bond in Cain-like and Adephagia-like capsids.

We next hypothesized that the handful of conserved amino acids found in the Bobi-like (15199) phages that were involved in capsid stability would be conserved across all of Structural Group 2; these phages all had very similar major capsid protein HK97-folds and we predicted they would use similar mechanisms to maintain the stability of the capsid around the local 3-fold. However, analysis of the amino acid sequences of the major capsid proteins of Structural Group 2 revealed almost no conserved amino acid sequence identify. A single conserved aspartic acid (D122 in Bobi) is found in all of Structural Group 2, located at the top of the spine helix (C-terminal end). In Bobi (and the other Bobi-like phages that we structurally characterized) this aspartic acid makes a hydrogen bond with the N-arm of the same major capsid protein chain. The hydrogen bond donor amino acid in Bobi is serine (S29), which is common across the majority of Structural Group 2. Some lack the serine and have either lysine or alanine in its place. Ziko is one such phage, which has a lysine at the equivalent position. In Ziko, the conserved aspartic acid interacts with the protein backbone of the lysine in the N-arm in a structurally equivalent position to serine 29 in Bobi.

Due to the lack of amino acid sequence identity, we turned to the models we had derived from the cryo-EM maps of the phages. We examined the structurally conserved interactions across Structural Group 2, once again hypothesizing that the most critical of these interactions would be conserved. We overlaid the local 3-fold axis of all six of the Bobi-like (pham 15199) phages and Ogopogo (pham 57445). To better represent more members of Structural Group 2 in this analysis we also included Cozz, a close relative of Ogopogo and also part of the 57445 pham (Supporting Figure 11). Cozz was subjected to cryo-EM analysis and a sub-3 Å resolution map was obtained that allowed for *de novo* model building. This structural comparison showed that there were no salt bridges conserved between the different phages in Structural Group 2, nor was there a conserved hydrogen bond network. Each phage used a different pattern of bond formation between the major capsid proteins for intermolecular stability. We, therefore, conclude that what unifies a Structural Group is the capsid configuration and that the phages have evolved different bond networks to achieve the same final product.

## Discussion

### Capsid stability

Tailed phages have been found in a wide range of environmental habitats, ranging from relatively benign soil to hot springs and stomach acid^27^. In addition to these harsh external environments, the capsid is also under stress from the inside: the predicted pressure that the packaged dsDNA exerts on the inside of the capsid has been estimated at 30 atmospheres^79^. The phage capsid must be stable enough to survive these two main stresses. Many stabilization mechanisms have been characterized at the local three-fold axis of the capsid, implying they play an important role. These typically include many inter-capsomer interactions, for example, the interaction of the P-domains around the three-fold axis of the capsid; the N-arm reaching across to make contacts with adjacent major capsid proteins, and the E-loop interacting with the P-domain. Only HK97 has had its isopeptide bond structurally characterized, although phages have been shown to use the isopeptide bond biochemically. Since then, each structurally characterized phage has been found to lack the isopeptide bond but uses alternate mechanisms to compensate for the lack of this bond to stabilize the local three-fold axis. These include extra domains in the HK97-fold, for example, the I domains found in P22^39^ and T4^48^, that have been shown to play a role in capsid stability^80^. Additional capsid proteins have also been characterized that are thought to play a role in stability. This includes minor capsid proteins/cement proteins found in several tailed phages and which form trimers/dimers throughout the capsid between the hexamers and pentamers. Other proteins, called decoration or ancillary proteins, have also been characterized that may be involved in stability, although in many cases they are not vital for capsid viability (for example, the soc protein of T4^81^). Finally, more diverse mechanisms have been characterized such as the lasso-like interactions in the E-loop observed in two phages isolated in hot springs^40, 41^.

However, some phages, for example, T7^26^ and the recently structurally characterized phage phiRSA1^27^, show that some phages rely solely on the electrostatic and hydrophobic interactions between the major capsid proteins^27^. Here, we have described a similar lack of stabilizing mechanisms in the T=9 actinobacteriophage Che8, which is an even more simplified example of the HK97-fold than phiRSA1; Che8 lacks the isopeptide bond and any other previously characterized capsid stabilizing interactions, instead relying on only a handful of protein: protein interactions between the major capsid proteins that are found in every HK97-fold major capsid protein. The Che8 major capsid protein also lacks the G-loop, which has been shown to play an important role in capsid assembly^82^. Furthermore, it lacks any potential loop that could compensate for the G-loop, demonstrating that the roles of the G-loop in the HK97 capsid are not required across all other HK97-folds. Additionally, Che8 has no minor capsid proteins, decoration proteins, or I-loops/other extended loops or embellishments that may contribute to capsid stability. This suggests that the core HK97-fold is all that is needed for capsid stabilization and that Che8 is likely to be more similar to the earliest HK97- fold. This is further supported by the Structural Group 9 phages that all have relatively small genomes (< 30 kbp) and are predicted to form T=4 or smaller capsids (unpublished data). All of these phages lack a G-loop and are similar in structure to the Che8 HK97-fold with a long spine helix. This leads us to speculate that the earliest common ancestor to these phages lacked the G-loop. It also suggests different assembly mechanisms between the different HK97-folds since the G-loop in HK97 was shown to play an important role in assembly and mutations in the G-loop led to the formation of aberrant particles^82^.

The isopeptide bond is a covalent bond between two neighboring major capsid protein subunits and is critical to the viability of HK97 virions. Here, we have structurally characterized other tailed phages that also use the isopeptide bond in their capsid. The Bobi-like (pham 15199) phages all use the same isopeptide bond as in HK97, although they substitute asparagine for aspartate in the P-domain to create the bond. The use of an aspartic acid to form the isopeptide bond has not been observed in the tailed phages before but has been characterized in bacterial proteins^83^. Also, the mechanism by which the isopeptide is formed may be subtly different. The catalytic glutamic acid residue is still present in the Bobi-like phages, but they lack two of the residues known to form the hydrophobic pocket that is important for the catalysis of the bond^24^. There are no obvious analogs in the Bobi-like phages for those two residues, and these phages may create the hydrophobic pocket through other means. However, not all of the Bobi-like phages use the isopeptide bond; a small subset of the Cluster K phages, which we term the Adephagia and Cain-like phages, do not use the isopeptide bond and the lysine in the E-loop is substituted with isoleucine. This resulting residue chemistry prevents the formation of an isopeptide bond. Phylogenetic analysis of the Bobi-like phages (Supporting Figure 8) suggests that the Cluster K phages diverged from within the Bobi-like phamily. Although this is speculative, it does support the model that the Adephagia- like and Cain-like phages had the isopeptide bond at some point before it was lost, as opposed to being an intermediate between non-isopeptide bond phages that then evolved to have the isopeptide bond. This is further supported by the other Cluster K phages having the correct lysine for isopeptide formation and presumably forming that isopeptide bond. We were unable to identify any unique increase in inter-capsid interactions in the Adephagia- and Cain-like phages that would compensate for the loss of the isopeptide bond. This suggests that, at least in the Bobi-like phages, the isopeptide bond is not critical to the viability of the phage capsid and that compensatory mechanisms, for example, minor capsid proteins, are not needed. This raises the question as to the role of the isopeptide bond, and why some phages do not require the extra stabilization it affords. A potential explanation is that Cain- and Adephagia-like phages package less dsDNA, exerting less internal pressure on the capsid than those that use the isopeptide bond. However, this correlation cannot yet be made as the amount of dsDNA packaged has not been measured, although we observe that both phages have cos-type genome ends that typically means that the DNA packaged is the same as the genome length.

### Capsid size

The tailed phages make protein shells of variable sizes. The smallest to date are the T=4 capsids of P68^30^ and the T=3 prolate phi29^28^. The majority are predicted to be T=7, although many “jumbo” phages have been characterized with very large T numbers^32^.

How capsid size is controlled is still an open question. However, many major capsid protein mutants that change the size of the final capsid have been identified in the model phages P22 and T4. The major capsid protein mutants in P22, where the capsid protein is referred to as the coat protein, all result in the wild-type T=7 capsid with the ability of also creating smaller T=4 capsids or aberrant particles^84^. Within the prolate phage T4, the mutants result in “giant” capsids that have lost the ability to regulate the length of the prolate caps and form very long prolate heads^85^. The work on *Staphylococcus aureus* infecting phages and the mechanisms that this bacterium uses to co-opt the phage capsids for the use of the bacteria all result in smaller capsids^34, 45^. Here we have identified several closely related phages that use related major capsid proteins from that same protein phamily, but make either T=7 or T=9 capsids. There are no obvious differences in structure or amino acid conservation between these T=9 and T=7 phage capsids (from phamily 15199) that explains the difference in size. The T=7 capsids use a different phamily of scaffold proteins than the T=9. supporting the role of the different scaffolding proteins as the main mechanism of capsid size determination. However, further work is needed to characterize the mechanisms by which these phages assemble. The actinobacteriophages are a rich resource for these types of studies. For example, the structural Group 1 phages contain both Che8 (a T=9 capsid) and Myrna, a T=16 capsid that uses minor capsid proteins^51^. Further study of the Group 1 phages could provide important insights into how minor capsid proteins are first incorporated into the capsid and how larger capsids evolve.

### The evolution of the major capsid proteins of the actinobacteriophages

We have reported here the first systematic structural study of a group of phages that infect hosts from a single phylum of bacteria. The actinobacteriophages infect hosts from an important bacterial phylum, with *Mycobacterium tuberculosis* a major health concern around the world and non-tuberculosis mycobacteria a major source of lung disease. Most phages are specific to a single host, although some can infect and replicate within many species of the same phylum. There are likely several reasons for the limited host range, including specificity of host binding proteins in phages, codon usage, differences in cellular machinery, and presence of phage-resistance mechanisms^86^. Phages are very likely to be limited to a specific phylum and to our knowledge presently there are no examples of phages that infect hosts in multiple bacterial phyla. The Actinobacteria are an ancient bacterial phylum and are estimated to have diverged from the Proteobacteria phylum over 3 billion years ago^87^. The majority of the well-characterized model system phages infect hosts in the Proteobacteria phylum, for example, P22 that infects *Salmonella* and T4 that infects *E.coli*, and most of the structures of tailed phages come from the Proteobacteria. The temporal distance between the Actinobacteria and Proteobacteria phyla, as well as the lack of evidence of phages moving between phyla, would suggest that the phages that replicate in the two phyla should also be temporally distant and that major capsid proteins will have had different evolutionary pathways. However, the structural analysis of the actinobacteriophages does provide evidence that these phages are ancient. The isopeptide bond in the Bobi-like phages provides some evidence that these HK97-folds predate the divergence of the Proteobacteria and Actinobacteria. Interestingly, it appears that structurally-related major capsid proteins can be found in different capsid morphologies. For example, Structural Group 1 from the actinobacteriophage structural dendrogram contains both the *Siphoviridae* Che8 and *Myoviridae* Myrna, both of which we have structurally characterized in this paper and previously^51^. How these two morphologies ended up using the same major capsid protein is intriguing, and additional characterization of Group 1 is needed to explore this further. Is there a T=9 *Myoviridae* in Group 1 that has the same capsid protein structure as Che8? We expect that further research into the different structural groups will provide a deeper insight into the evolutionary relationship between tailed phages.

## Supporting information

Alphafold PDB files and PDB validation files

## Acknowledgments

We thank Dr. Gabrielle Valles for a helpful review of the paper. We acknowledge the hard work and dedication of all those involved (those at the University of Pittsburgh and HHMI) in the creation and continued support of the SEA-PHAGES program.

Specifically, we thank the following students from the SEA-PHAGES program for the isolation of each phage:

Bridgette: Kira Zack and others at the University of Pittsburgh, PHIRE Program

Muddy: Lilli Hoist and others at the University of Kwazulu-Natal, PHIRE Program

Oxtober96: Lijia Xin and others are the University of Connecticut, SEA-PHAGES Program

Ziko: Anna Bondonese and others are at the University of Pittsburgh, SEA-PHAGES Program

Bobi: Margaret Korty and Stephanie Maas and others are Purdue University, SEA-PHAGES Program

Adephagia: Jordan L. Mosier and others at the University of North Texas, SEA-PHAGES Program

Cain: Thomas Cast and Kara Gallo and others at Gonzaga University, SEA-PHAGES Program

Che8: V. Kumar and others at the Albert Einstein College of Medicine

Ogopogo: Kaylee Nicholson and others at the University of California, Santa Cruz, SEA-PHAGES Program

Cozz: Matthew Montgomery and others at the University of Pittsburgh, PHIRE Program

We also want to thank the myriad of students and faculty who have contributed to the 201 phages we included in our bioinformatic analyses.

A portion of this research was supported by NIH grant U24GM129547 and performed at the PNCC at OHSU and accessed through EMSL (grid.436923.9), a DOE Office of Science User Facility sponsored by the Office of Biological and Environmental Research.

This work was supported by National Institutes of Health grants GM131729 and Howard Hughes Medical Institute grants GT12053 (to GFH). The University of Pittsburgh Titan Krios microscope and Falcon 3 camera were supported by the Office of the Director, National Institutes of Health, under award numbers S10 OD025009 and S10 OD019995, respectively (JFC).

We also thank the following scientists at PNCC for the data collection: Theo Humphreys, Omar Davulcu, Nancy Meyer, and Rose Marie Haynes.

## Supporting information

**Supporting Table 1.** Host and bacteriophage information for each bacteriophage used in this study.

| Phage name | MCP<br>pham | Cluster/Sub-<br>cluster | Host | Genome<br>length | Growth<br>temp.<br>(°C) |
| --- | --- | --- | --- | --- | --- |
| *Oxtober96* | 15199 | EA/EA1 | *Microbacterium*<br>*foliorum* NRRL B-24224 | 41798 | 32 |
| *Bridgette* | 15199 | FA | *Arthrobacter globiformis* B-2979 | 43113 | 32 |
| *Muddy* | 15199 | AB | *Mycobacterium*<br>*smegmatis mc<sup>2</sup>155* | 48228 | 37 |
| *Bobi* | 15199 | F/F1 | *Mycobacterium*<br>*smegmatis mc<sup>2</sup>155* | 59179 | 37 |
| *Adephagia* | 15199 | K/K1 | *Mycobacterium*<br>*smegmatis mc<sup>2</sup>155* | 59646 | 37 |
| *Cain* | 15199 | K/K6 | *Mycobacterium*<br>*smegmatis mc<sup>2</sup>155* | 60813 | 22 |
| *Ziko* | 15199 | DP | *Gordonia terrae* 3612 | 68860 | 32 |
| *Ogopogo* | 57445 | F/F1 | *Mycobacterium*<br>*smegmatis mc<sup>2</sup>155* | 56867 | 37 |
| *Cozz* | 57445 | CT | *Gordonia terrae* 3612 | 46600 | 32 |
| *Che8* | 4631 | F/F1 | *Mycobacterium*<br>*smegmatis mc<sup>2</sup>155* | 59471 | 37 |

**Supporting Table 2.**
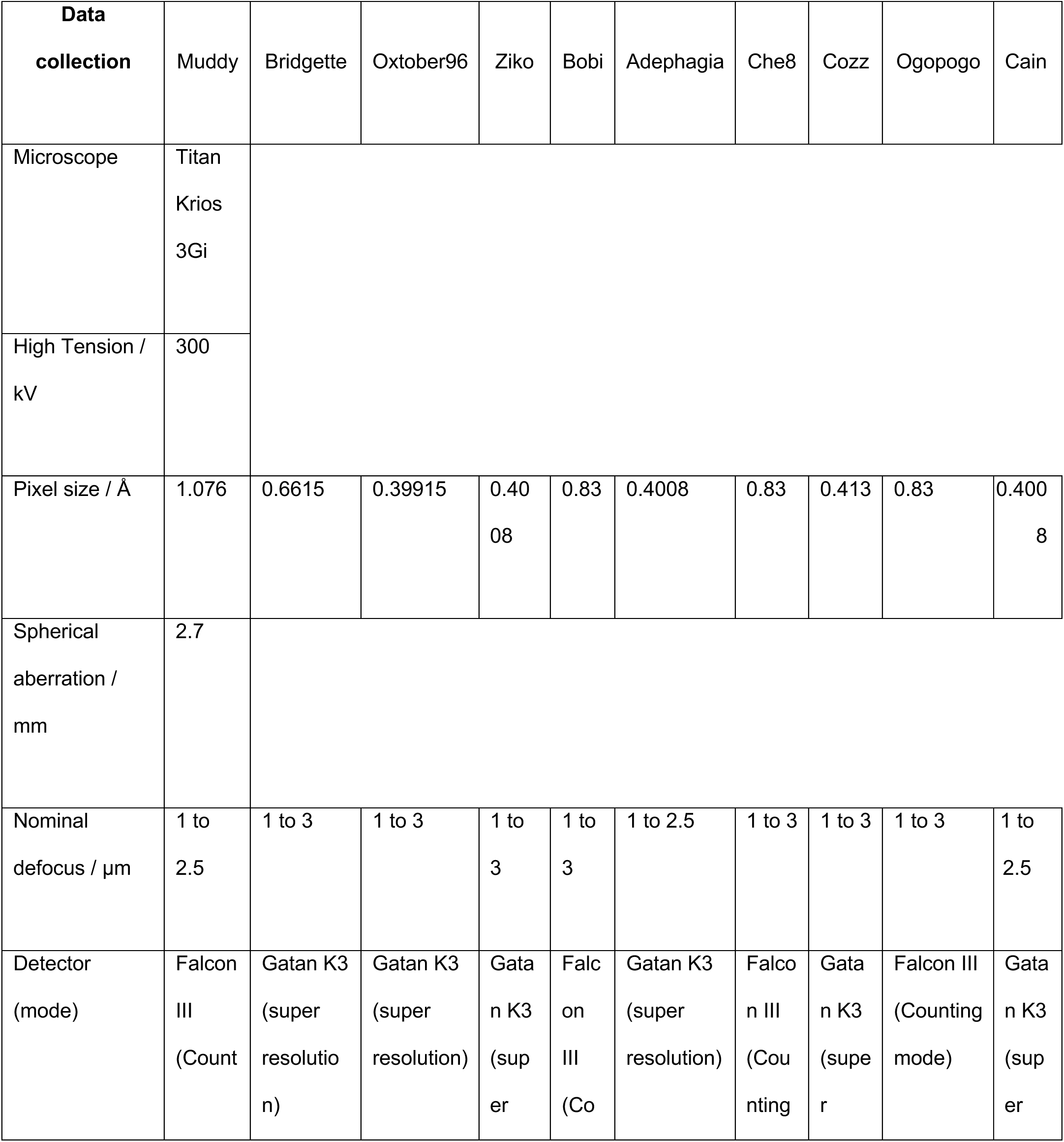
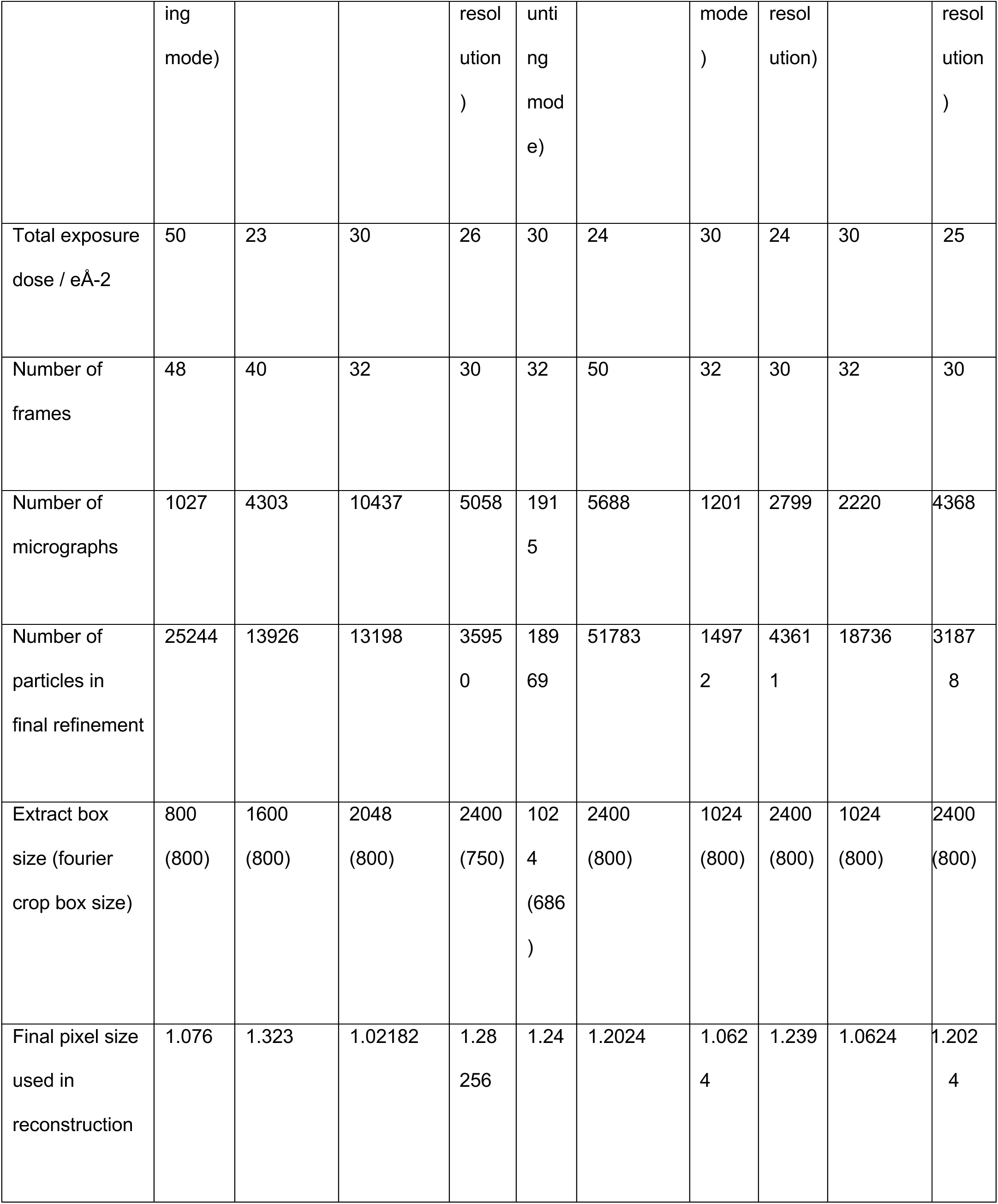
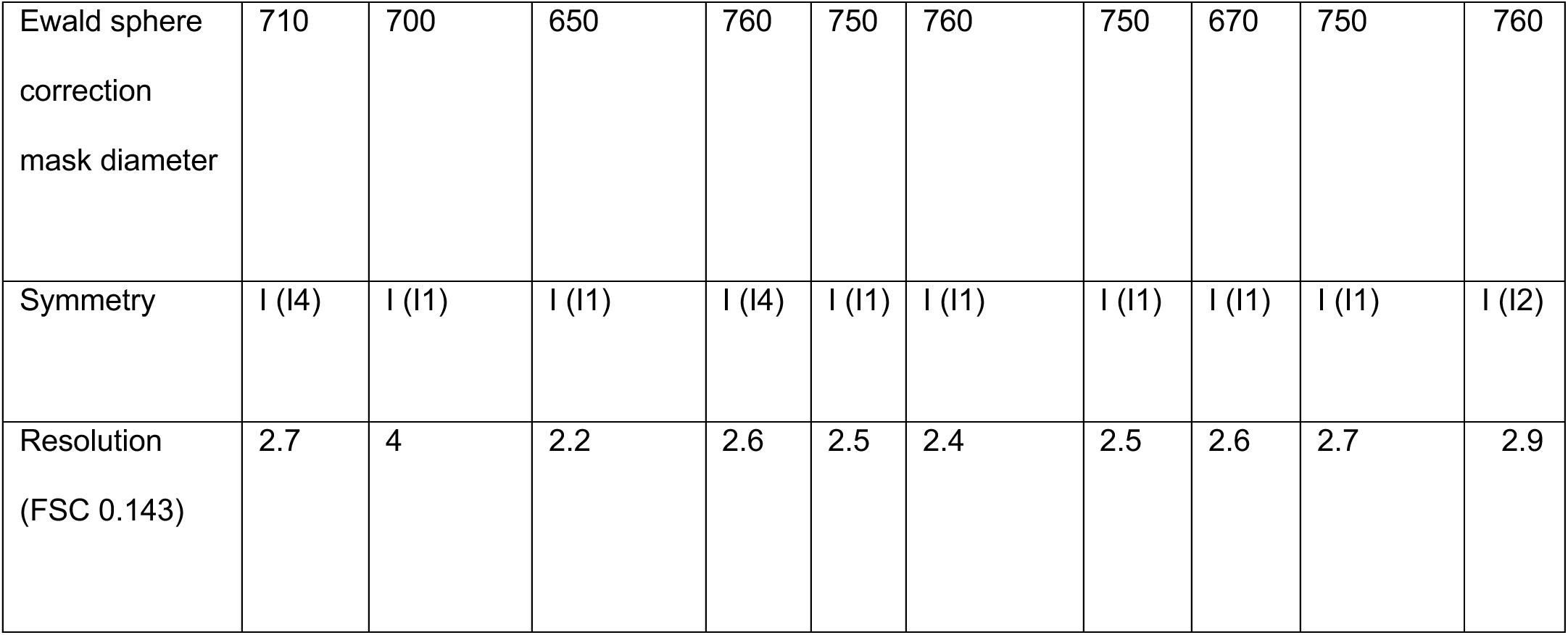
Cryo-EM collection parameters, analysis, and final resolutions.

**Supporting Table 3.** RMSD values of the Bobi-like (pham 15199) cryo-EM derived major capsid protein models when compared to Bobi. Values are calculated using the Matchmaker command in ChimeraX with default settings.

|  |  |
| --- | --- |
| *Muddy* | 1.0 |
| *Bridgette* | 1.2 |
| *Oxtober96* | 1.1 |
| *Ziko* | 1.1 |
| *Adephagia* | 0.6 |

**Supporting Figure 1.**
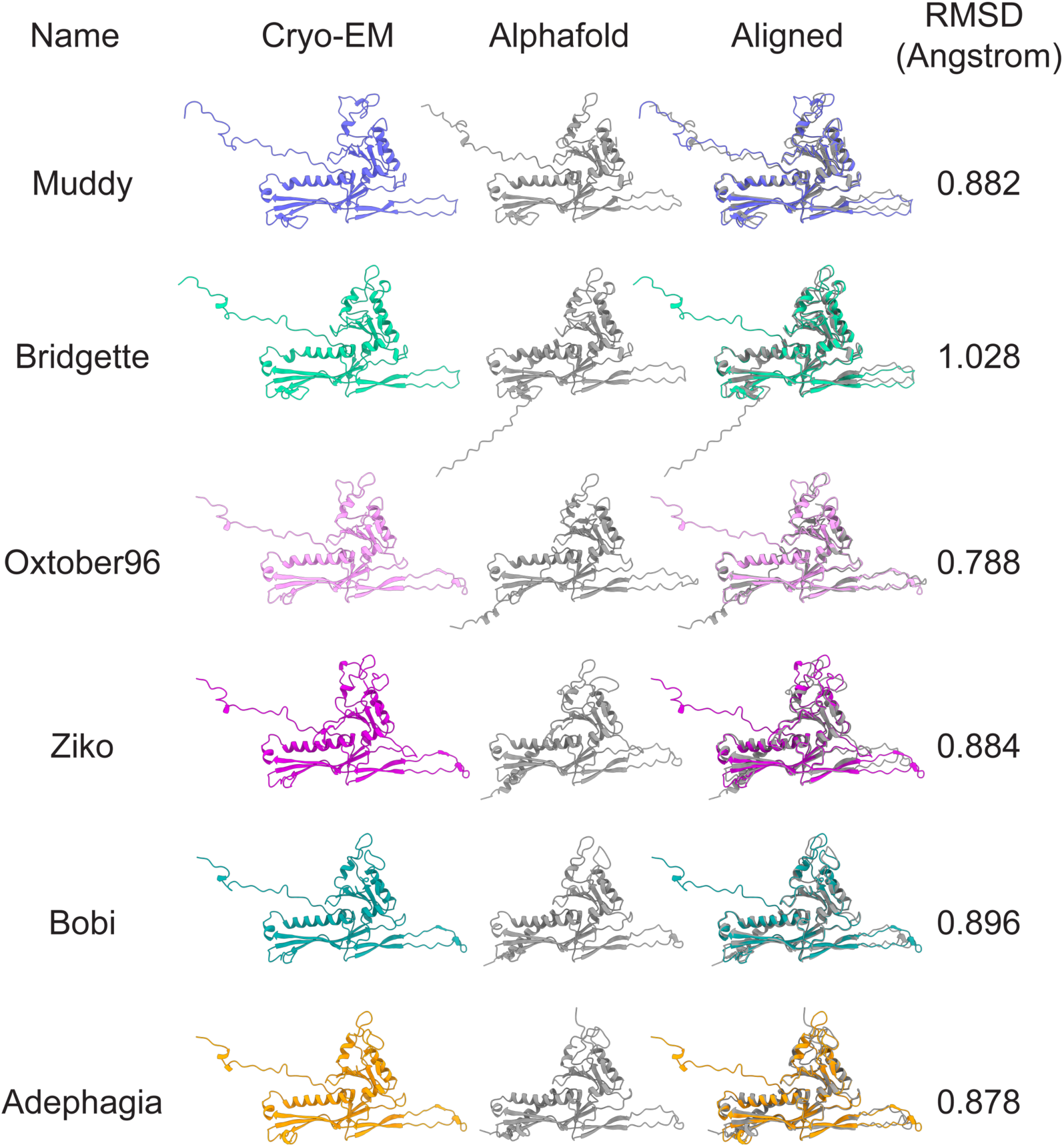
Comparison of cryo-EM derived models and Alphafold predictions of the six major capsid proteins of the Bobi-like phages (pham 15199).

**Supporting Figure 2.**
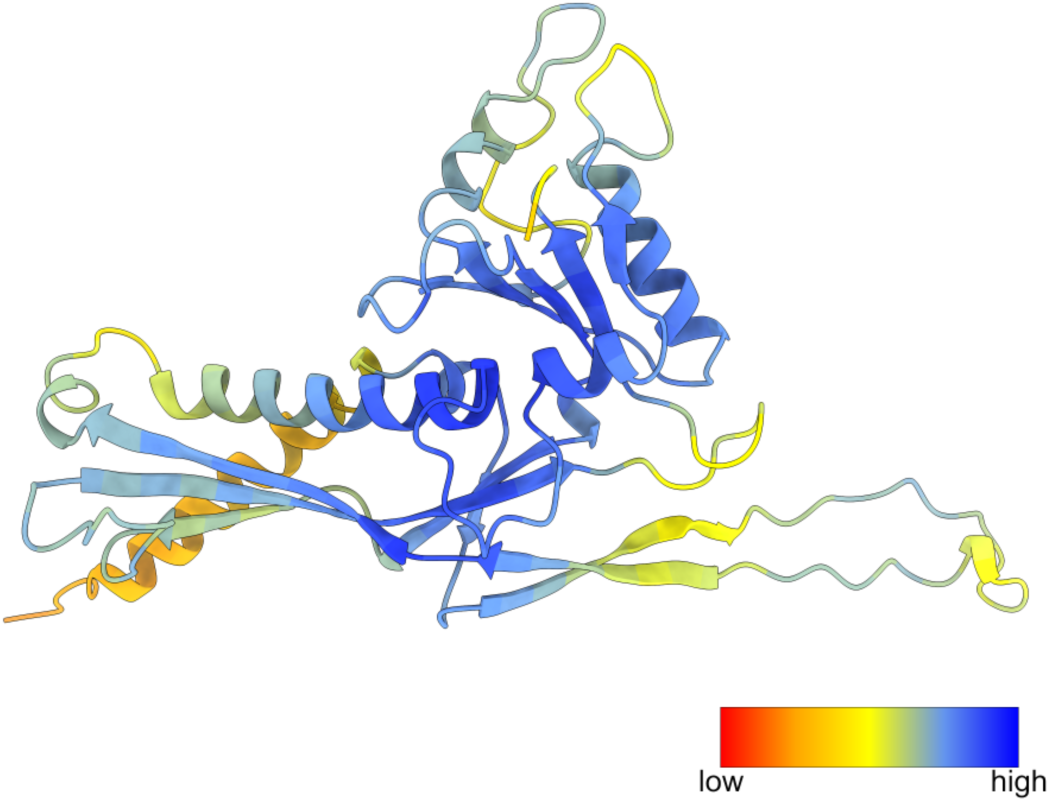
Confidence levels of Bobi major capsid protein predicted by Alphafold. The level of confidence is colored from low (red) to high (blue).

**Supporting Figure 3.**
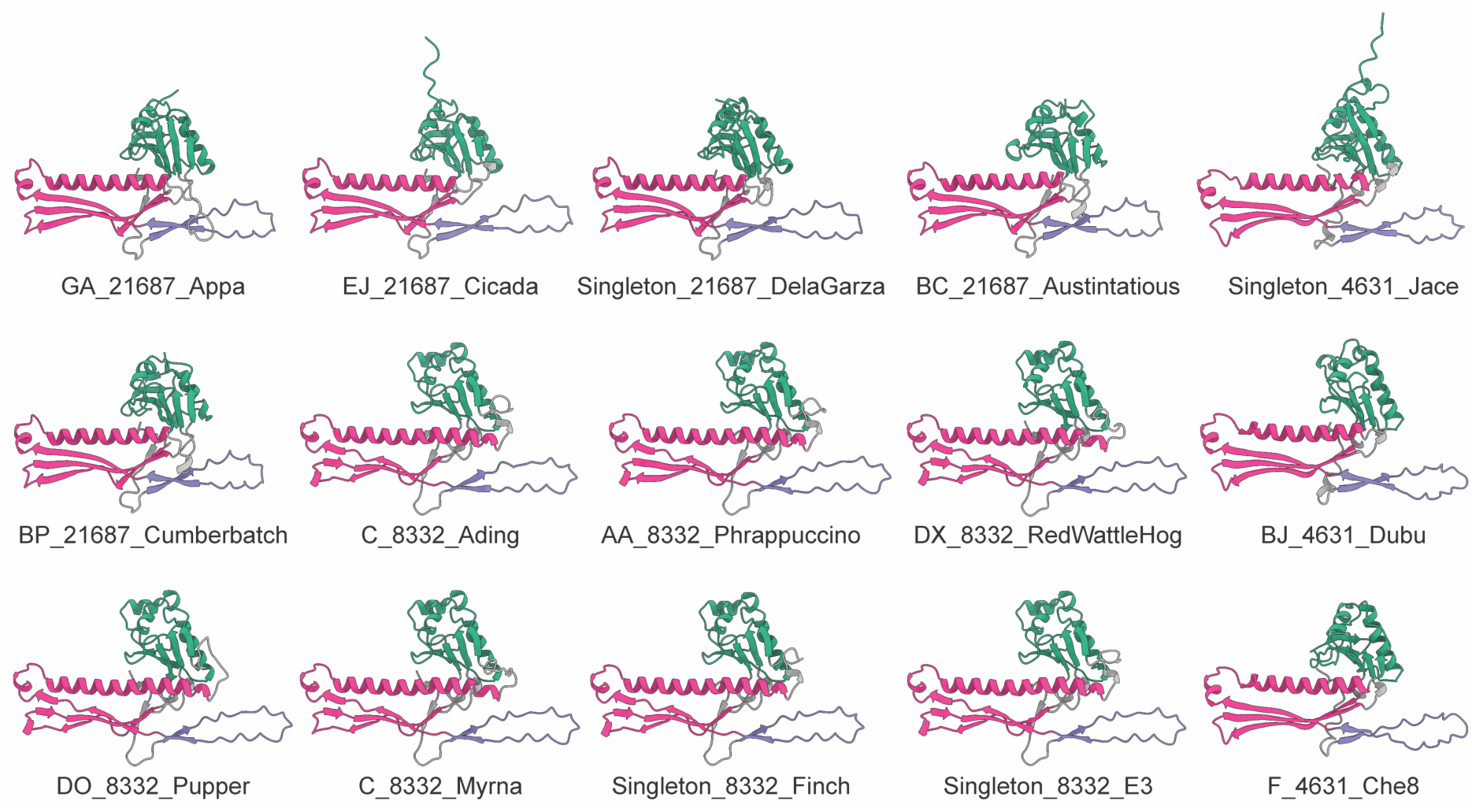
Alphafold predictions of the major capsid proteins from structural Group 1. All major capsid proteins had their N-terminal removed as described in the text. The full-length and truncated PDB files of the predicted structure can be found in Supporting Files.

**Supporting Figure 4.**
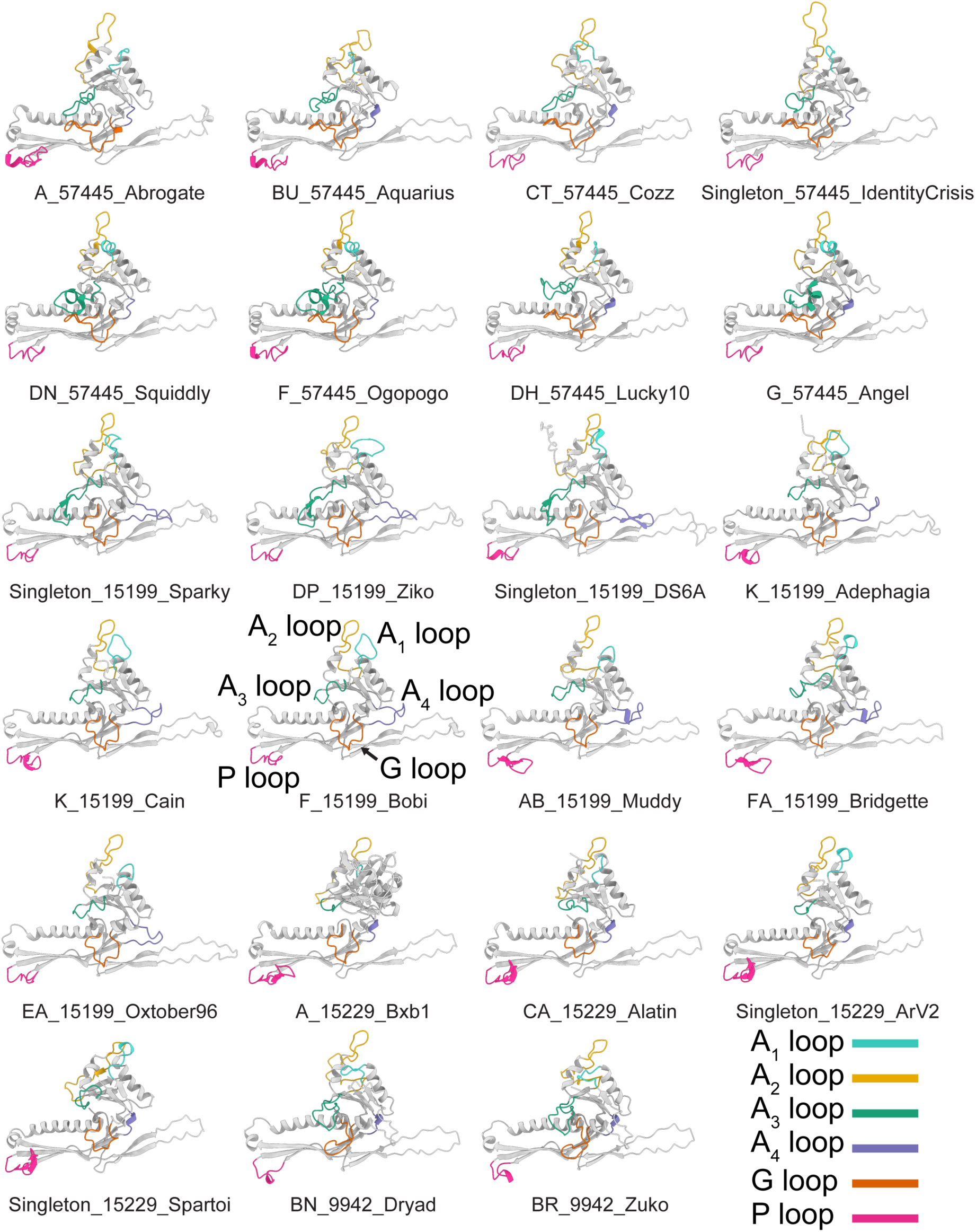
Alphafold predictions of the major capsid proteins from structural Group 2. All major capsid proteins had their N-terminal removed as described in the text. The full-length and truncated PDB files of the predicted structure can be found in Supporting Files.

**Supporting Figure 5.**
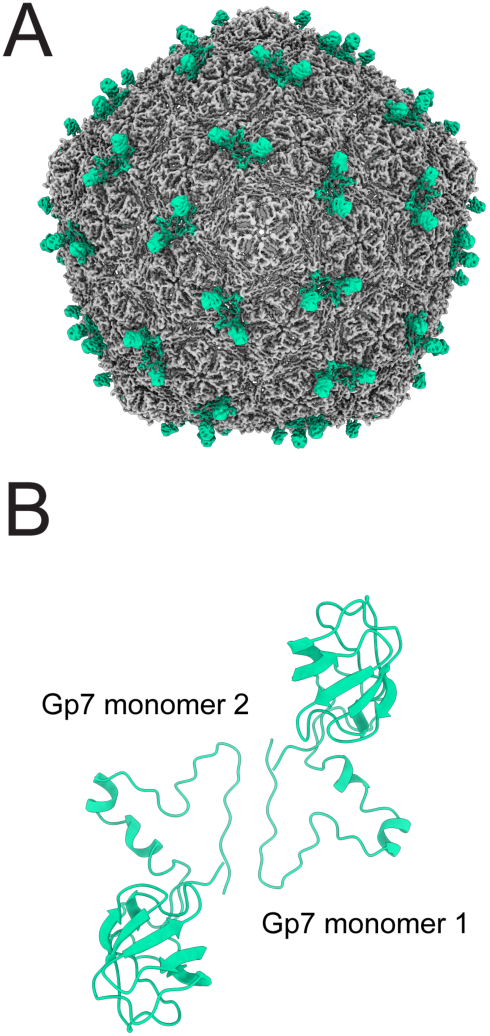
The decoration protein of Bridgette (Gp7). A shows the entire Bridgette capsid with the decoration protein dimers (Gp7) colored. B shows the model of the Gp7 (amino acids 2-125) dimer in the context of the capsid.

**Supporting Figure 6.**
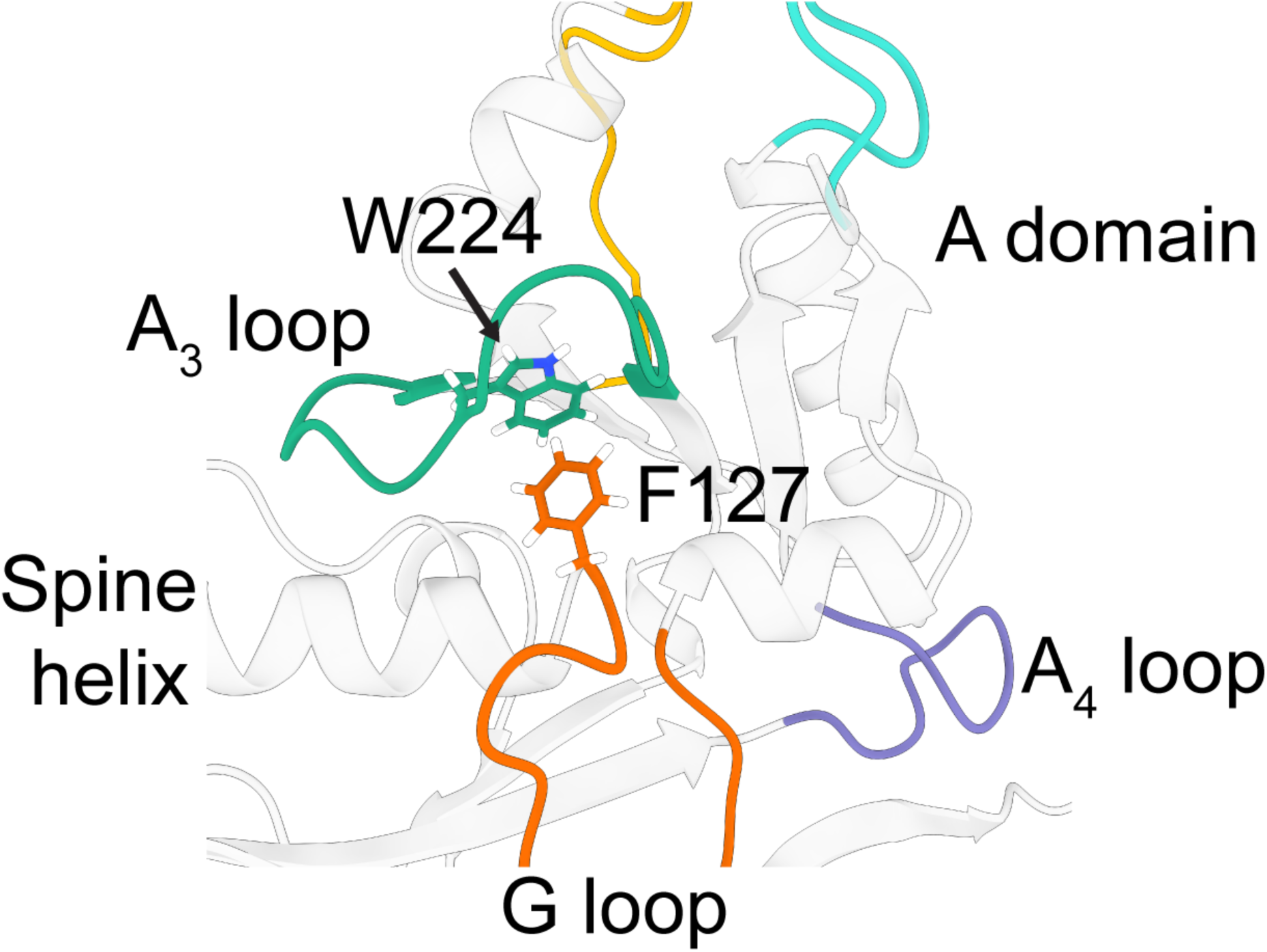
Pi-pi interaction within the Bobi major capsid protein. Tryptophan 224 in the A3 loop forms a pi-pi interaction with Phenylalanine 127 in the G-loop.

**Supporting Figure 7.**
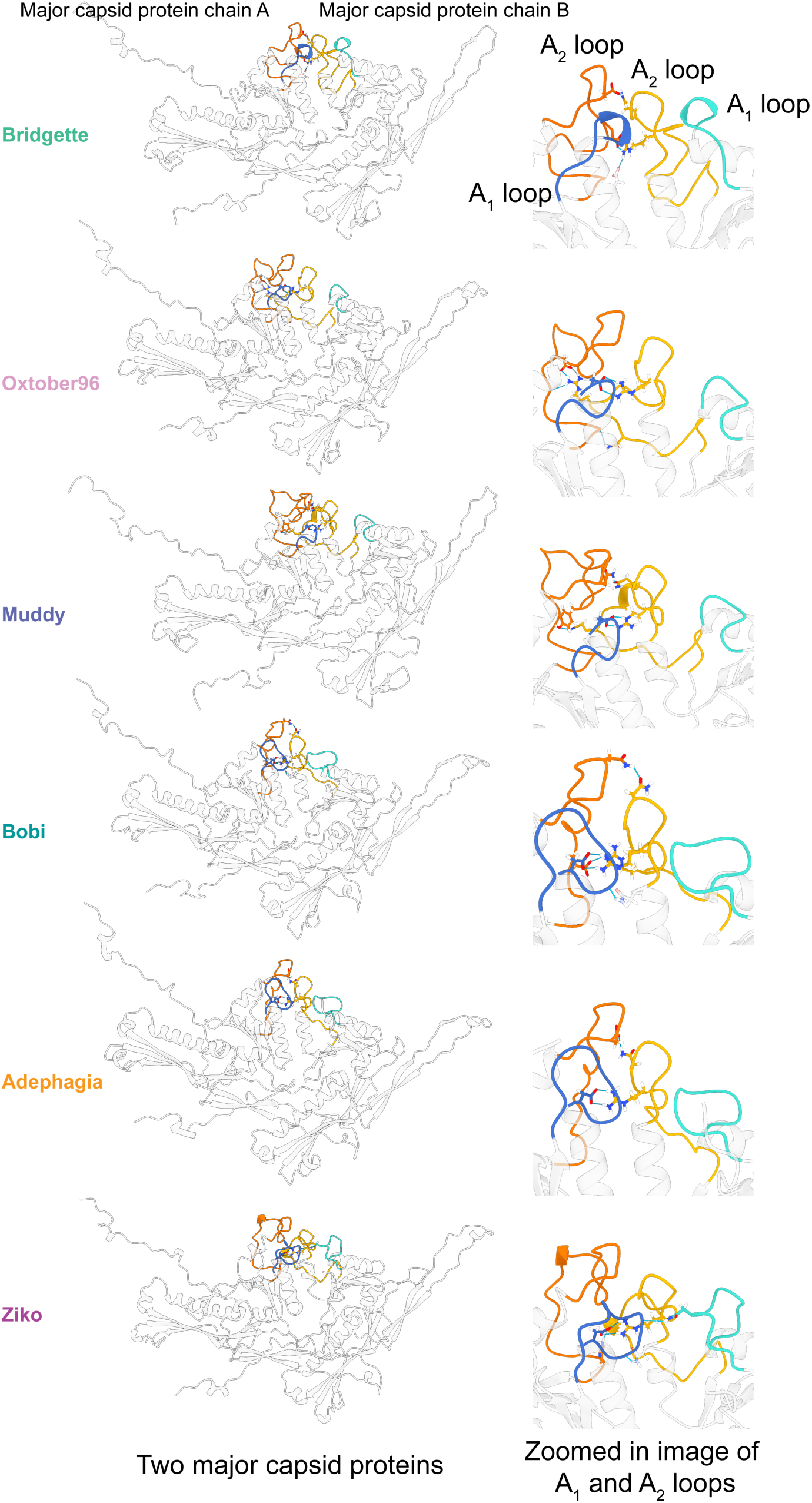
A_1_ and A_2_ loop interactions of the Bobi-like (15199) bacteriophages.

**Supporting Figure 8.**
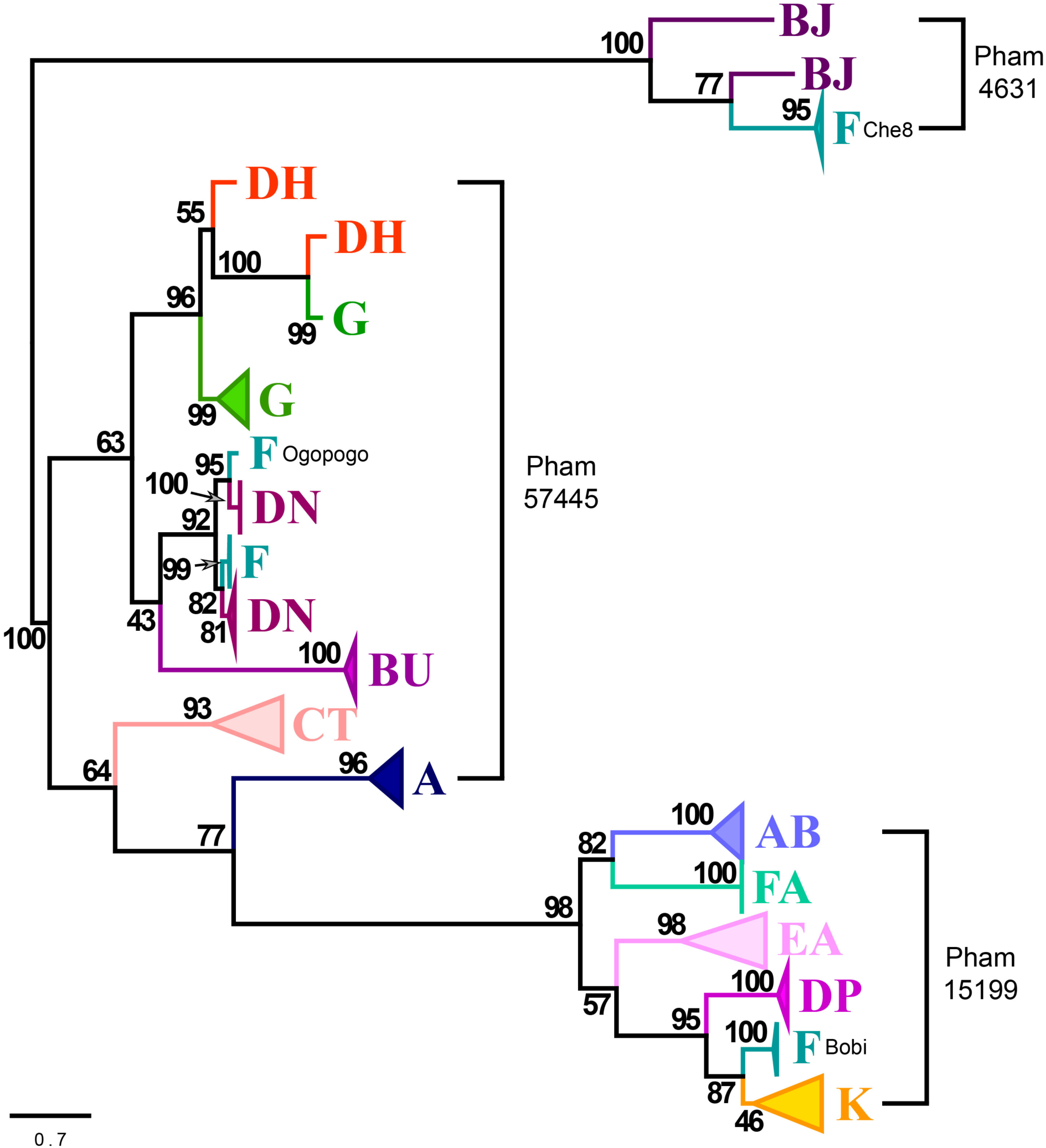
Unrooted phylogenetic tree of the 722 major capsid proteins of the Che8-like bacteriophages (pham 4631); the Ogopogo-like bacteriophages (pham 57445) and the Bobi-like bacteriophages (pham 15199). The tree has been collapsed for clarity. A pdf of the full un-collapsed tree can be found in the Supporting files.

**Supporting Figure 9.**
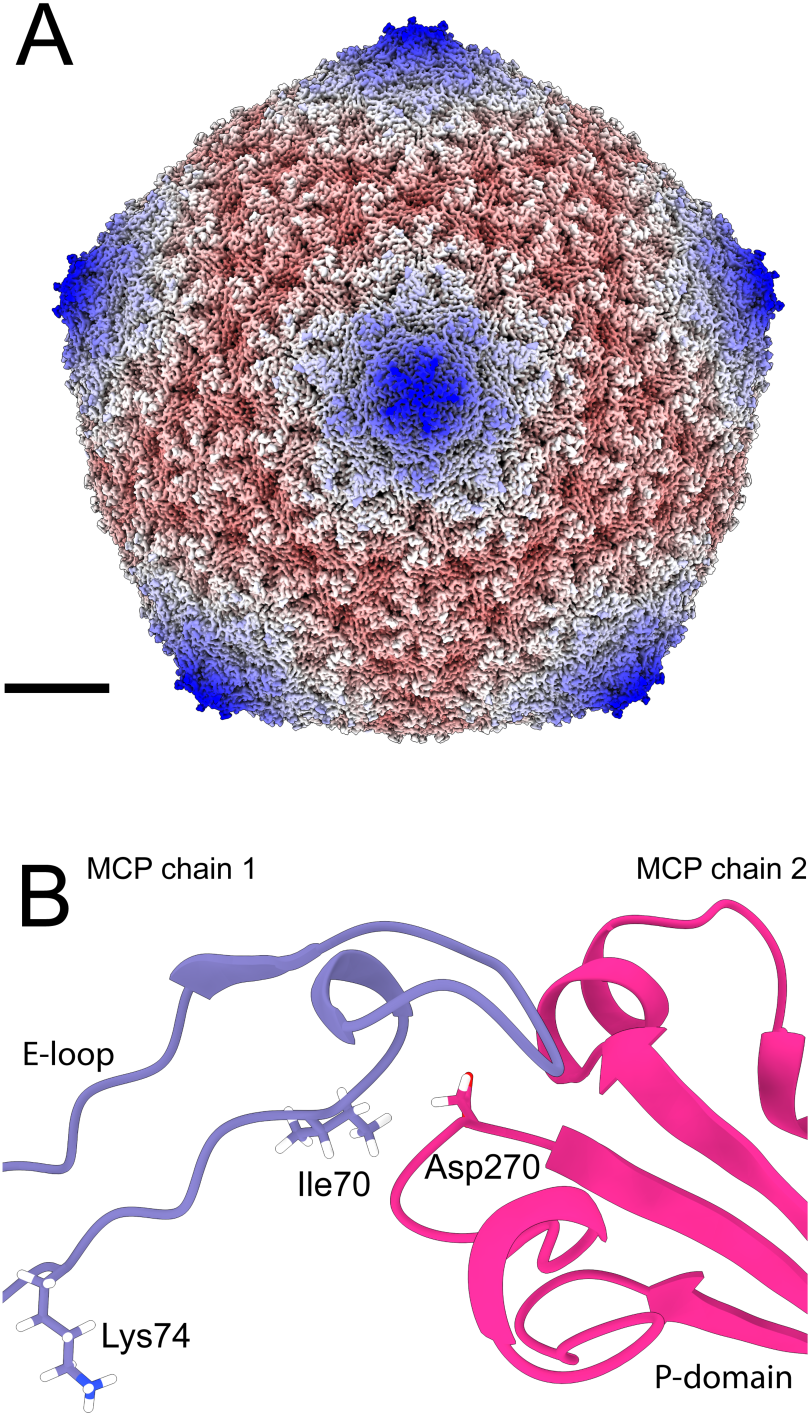
Cain (K cluster, 15199 MCP pham) does not have an isopeptide bond. A) Map of Cain, colored by radius using the same color scheme as in Figure 4. B) Image of two major capsid proteins of Cain around the 3-fold axis highlighting Asp270 and Ile70 that are in equivalent positions of the amino acids in Bobi that forms the isopeptide bond. Lys74 is also highlighted to show it is too distant to be part of an isopeptide bond with the P domain of MCP chain 2.

**Supporting Figure 10.**
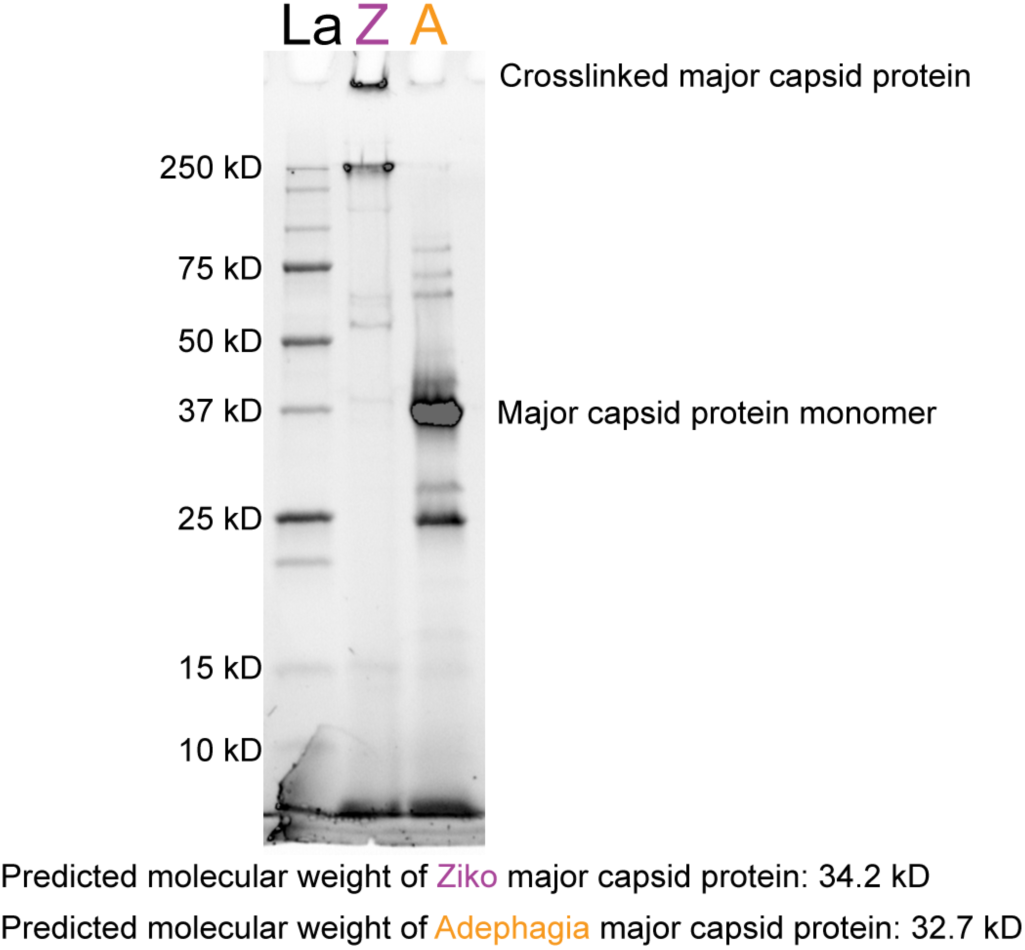
SDS-PAGE of Ziko (Z) and Adephagia (A) bacteriophages. Cesium chloride purified capsids of Ziko and Adephagia were diluted to 1 mg/mL and mixed with Laemmli buffer and boiled for 10 minutes at 95°C. Boiled samples were loaded onto a 10% SDS-PAGE Bio-Rad stain-free TGX gel and run for 1 hour at 180 V. Protein bands were visualized using the stain-free TGX technology. Ladder (La) was Unstained Precision Plus Protein Standard from Bio-Rad.

**Supporting Figure 11.**
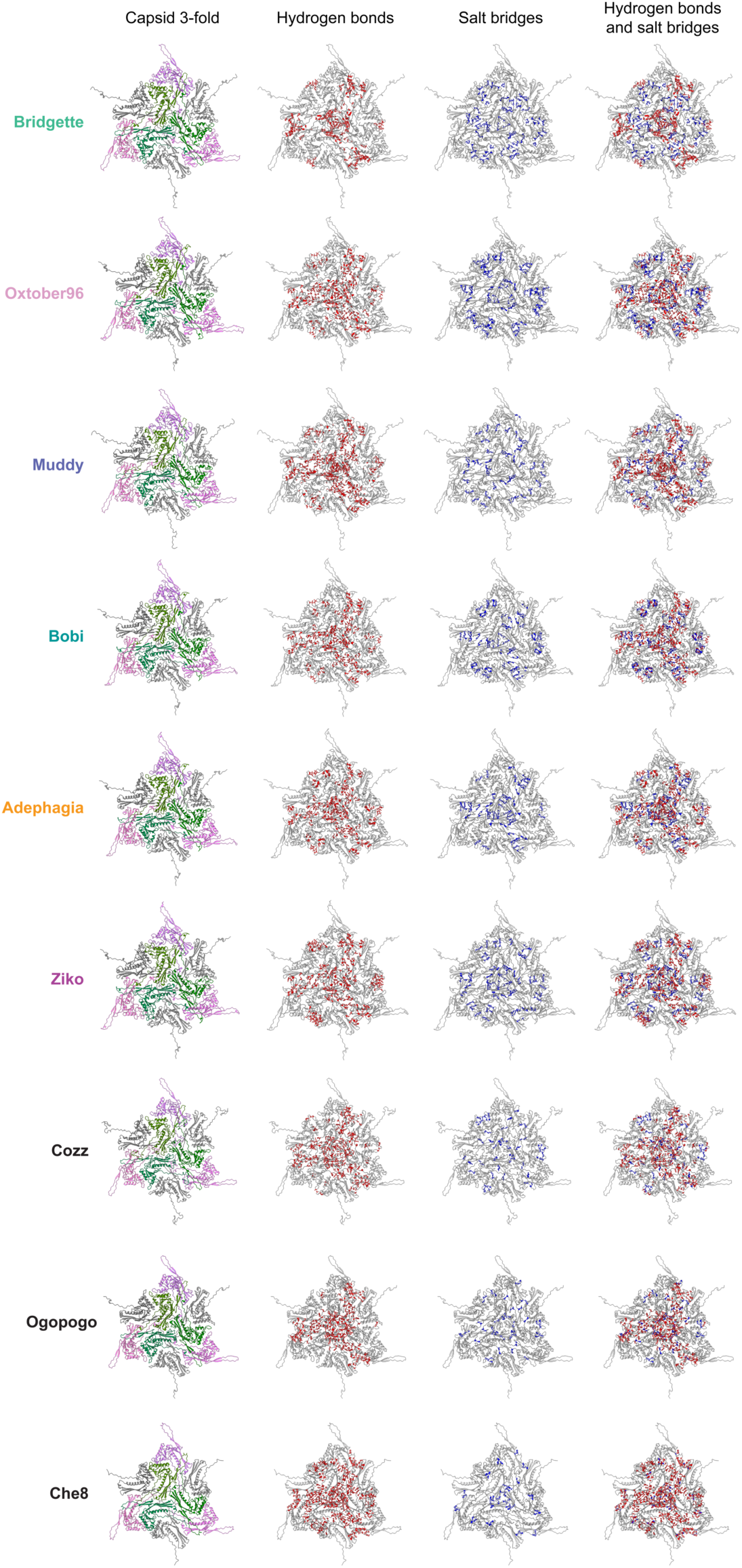
The three-fold axes salt bridges and hydrogen bond distribution around the 3-fold axes of the capsid. The nine major capsid proteins interacting around the three-fold axes are shown for each of the Bobi-like (15199) bacteriophage capsids. The hydrogen bond (red) and salt bridge (blue) networks are shown separately and overlaid.

